# Eye and hand coarticulation during problem solving reveals hierarchically organized planning

**DOI:** 10.1101/2024.11.18.624090

**Authors:** Mattia Eluchans, Antonella Maselli, Gian Luca Lancia, Giovanni Pezzulo

## Abstract

During everyday activities—such as preparing a cup of coffee or traveling across cities—we often plan ahead and execute sequences of actions. However, much remains to be understood about how we plan and coordinate such sequences (e.g., eye and hand movements) to solve novel and challenging tasks, for which plans must be formed from scratch. This study investigates how participants coordinate gaze and cursor movements during problem solving tasks that involve selecting a trajectory on a grid connecting multiple targets. By focusing on the action execution phase, we aimed to probe the structure of the gaze-cursor plans that participants used to solve the tasks. Our analysis reveals three main findings. First, consistent with previous studies, participants segment the problem into sequences of gestures; within each gesture, gaze focuses on a target and remains fixed until the cursor reaches it, then shifts to the next target. Second, both gaze position—while fixating on the current target—and the kinematics of cursor movement leading up to the current target allow prediction of the next cursor movement’s direction, revealing coarticulation in both cursor-cursor and gaze-cursor movements. Third, and most interestingly, the position of the gaze around the current target aligns with the direction of the next saccade, revealing coarticulation between successive gaze fixations. Together, these findings show that participants break the problem into gesture sequences and plan multiple eye and cursor movements in advance to efficiently reach both the current and upcoming gesture targets. This suggests a hierarchical planning strategy, with participants planning ahead at two levels: gesture targets and cursor movements.

## Introduction

Most of our daily activities, such as preparing a cup of coffee, traveling to a destination, or setting up a table for dinner, require planning and executing sequences of actions that achieve the intended goals. Although human planning has been a central concern of cognitive science since its inception (Newell et al., 1972), much remains to be understood about how we plan and coordinate the sequences of actions (e.g., eye and hand movements) necessary to solve cognitive problems. Previous studies investigated human planning during the solution of various types of problems, such as when navigating mazes, solving games such as chess, or puzzles like the tower of Hanoi. Some studies used the time taken before making a choice as an index of planning complexity (Stone, 1960; Ratcliff, 1978; Fudenberg et al., 2019; Myers et al., 2022). Other studies used computational models to analyse participants’ sequential action selection (e.g., the route taken in a maze), revealing various approximations to optimal planning strategies that emerge due to limited temporal and cognitive constraints (Eluchans et al., 2024; Mattar and Lengyel, 2022; Ho et al., 2022; Tomov et al., 2020; Donnarumma et al., 2016; Velasco et al., 2025; Kessler et al., 2024; De Cothi et al., 2022).

Most of the above studies focus on planning sequences of abstract actions. However, in real-life situations, planning abstract action sequences often needs to be translated into the execution of coordinated movements. For example, making a cup of coffee requires not only identifying and ordering the correct preparation steps – such as adding water and coffee grounds, lighting the stove, and serving the coffee – but also translating these steps into sequences of coordinated eye and hand movements (Maselli et al., 2023; Lepora and Pezzulo, 2015; Cisek and Pastor-Bernier, 2014; Yoo et al., 2021; Reina, 2003; Spivey, 2006; Rothkopf et al., 2007; Hayhoe et al., 2012). The close relationship between abstract cognitive planning and real-time movement control raises the possibility of using movement kinematics as a window into planning processes. In keeping, one line of research investigates planning processes by analysing gaze behaviour before and during the execution of action sequences (Rayner, 2009; Zhu et al., 2022; Pelz and Canosa, 2001; Binder et al., 2023; Najemnik and Geisler, 2005; Baldauf and Deubel, 2008; Lakshminarasimhan et al., 2020; Kashefi et al., 2024; Arato et al., 2024; Grant and Spivey, 2003). These studies reveal that gaze behaviour is driven by both salient visual stimuli and (present and future) goals (Mannan et al., 1997; Itti and Koch, 2001; Parkhurst et al., 2002; Parkhurst and Niebur, 2003; Rothkopf et al., 2016; Hayhoe and Ballard, 2005; Hayhoe and Rothkopf, 2011; Zhao and Marquez, 2013; Adams et al., 2015). During preparatory phases before the execution of action sequences, gaze can be used to scan the environment and to gather information about the problem (Crowe et al., 2000; Fiedler and Glöckner, 2012; Baldauf et al., 2006; Baldauf and Deubel, 2010; Diamond et al., 2017; Todorov, 1998; Gordon et al., 2025). Subsequently, during motor execution, most gaze fixations are on objects immediately needed but some are on objects needed in the future (Pelz and Canosa, 2001; Baldauf, 2018). For example, gaze during a tic-tac-toe game could predict participants’ future moves, with accuracy increasing with their expertise (Huang et al., 2023). Further analysis of gaze behavior in route planning tasks reveals the presence of backward and forward planning (Zhu et al., 2022). Moreover, studies on attention allocation during saccade sequences reveal that saccades are planned simultaneously rather than sequentially during perceptual discrimination tasks, and their amplitude and direction are affected by subsequent saccades as part of instructed sequences (De Vries et al., 2014; Baldauf and Deubel, 2008; McPeek et al., 2000; Ames et al., 2019; Azadi et al., 2021; Robinson, 1963). Another line of research reveals that it is possible to identify sequential planning by analysing movement *coarticulation*, or the fact that when two movements in a sequence are planned, the execution of the first movement changes as a function of the second movement to be executed. For example, phonology studies assessed that the same letters are pronounced differently, depending on the next letters to be uttered (Whalen, 1990; Daniloff and Hammarberg, 1973; Fowler and Saltzman, 1993). Coarticulation has been reported in various other domains, such as in fingerspelling, motor skill performance and acquisition, suggesting its generality (Shah et al., 2013; Jerde et al., 2003; Maselli et al., 2025).

While the above studies show that movement kinematics provide information about covert planning, they mostly focused on familiar and simple tasks. Planning during novel and challenging problem solving tasks remain largely unaddressed. To address this challenge, we tracked and analysed participants’ gaze and mouse cursor kinematics while they solved novel planning tasks on a PC (see Figure 1 for an example).

**Figure 1:**
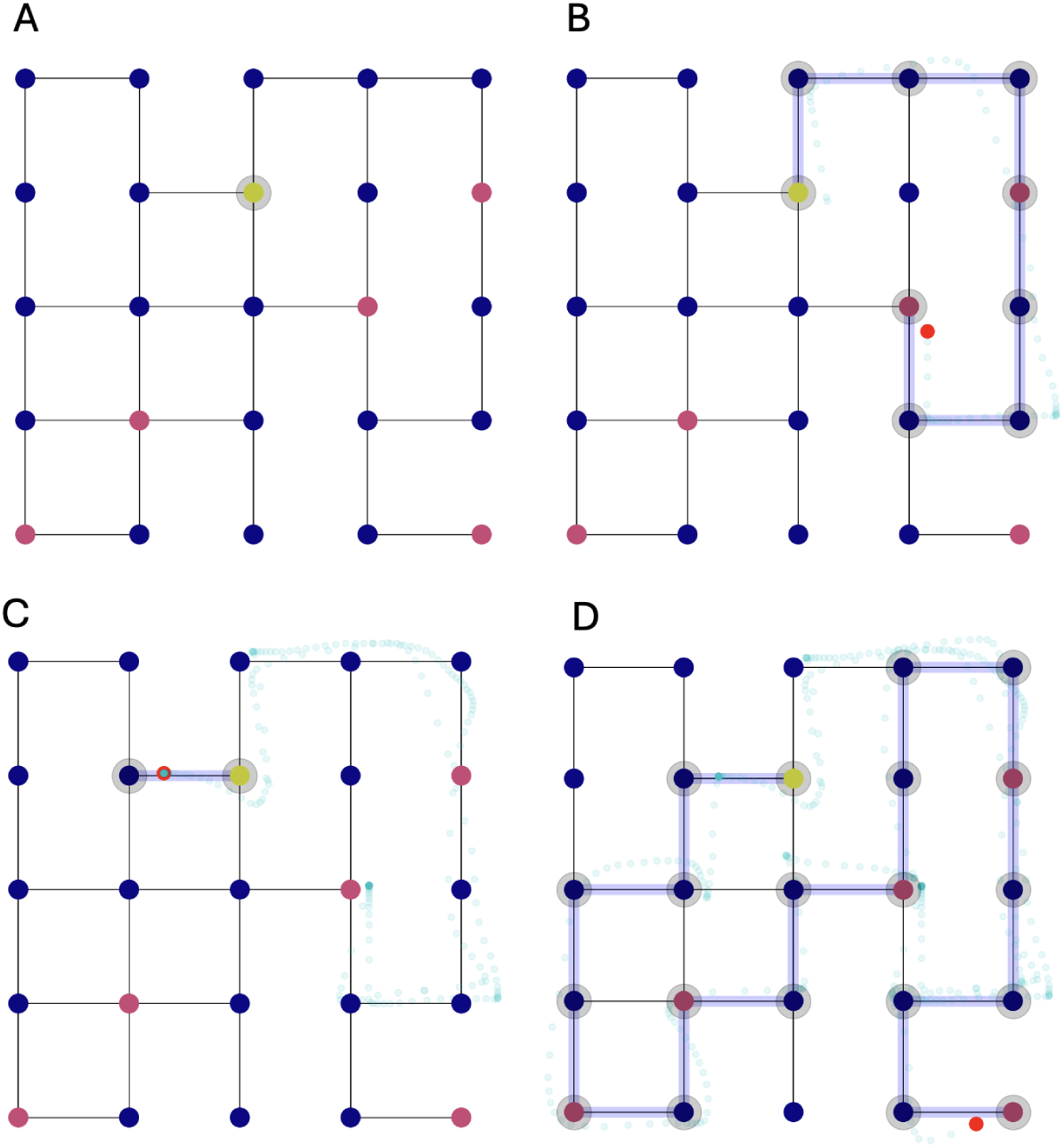
An example of a problem that participants faced in this study. The problem requires finding a path in the grid that starts from the start location (yellow node) and collects all the “gems” (red nodes), without passing twice through the same node. Participants solved the problem by controlling a computer mouse on a PC and dragging with a continuous movement the cursor over the graph: once the cursor entered a(n invisible) collider the node was darkened. The figure shows four time steps of the solution of an example participant. The blue path shows the path taken by a participant (which was visible to the participant), the azure dots show the cursor trajectory sampled at 60 Hz and the small red dots show gaze positions (both of which were not visible to the participant). The participant sees the grid (A) and chooses an incorrect path toward two gems on the right (B), then backtracks to the starting point (C) and finally identifies the correct path to solve the problem (D).

We aimed to address three main questions. First, we asked whether gaze and cursor dynamics reveal if and how participants segment a motor plan into sequences of movements, and with which properties. Second, we asked whether participants’ gaze position and/or cursor kinematics permit predicting the next cursor direction, therefore showing cursor-cursor and/or gaze-cursor coarticulation. Third, we asked whether gaze fixations provide information about the next saccade direction, therefore showing gaze-gaze coarticulation. Examples of cursor-cursor, gaze-cursor and gaze-gaze coarticulation (or a lack of it) in our task are shown in Figure 2.

**Figure 2:**
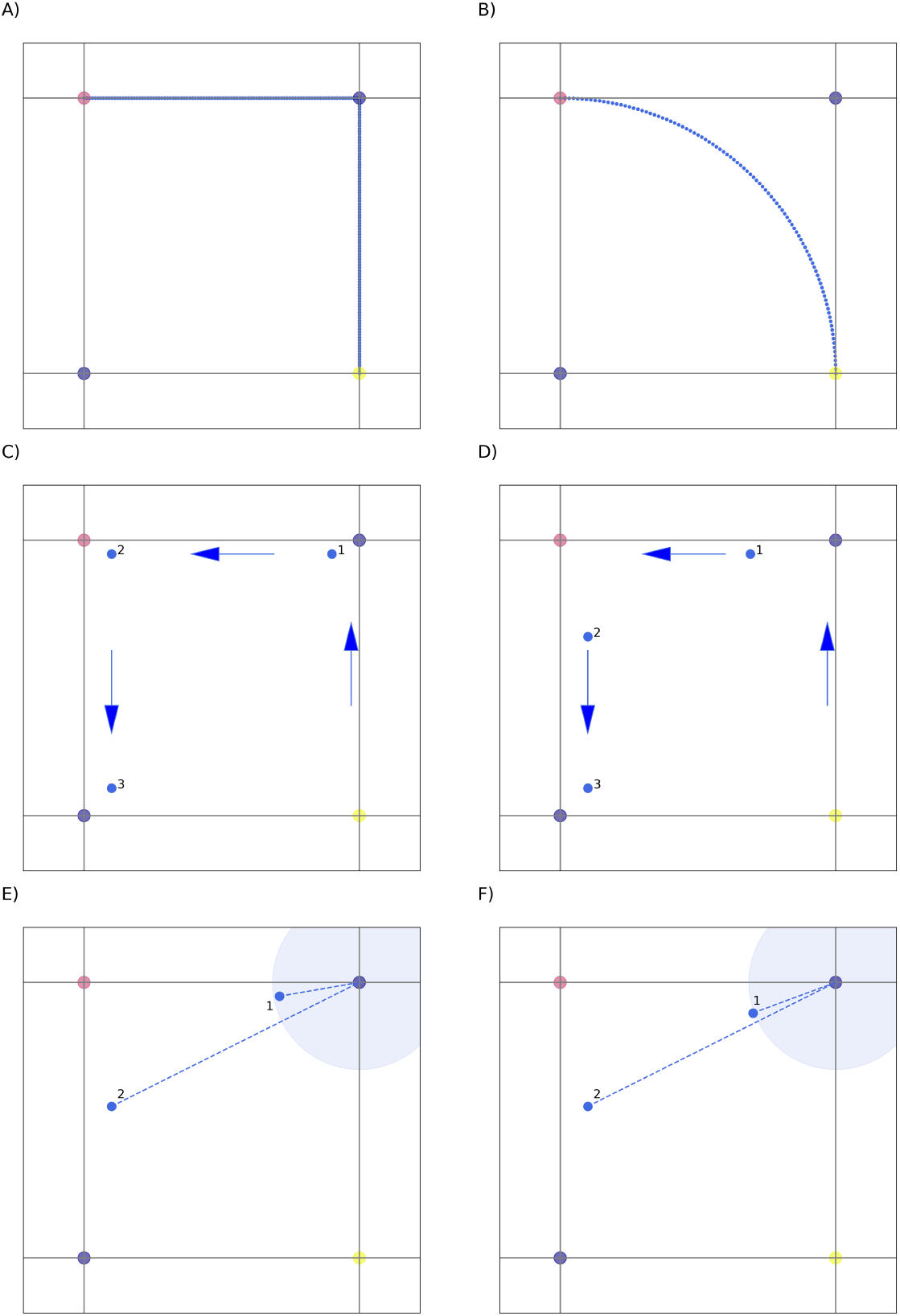
Examples of the three types of coarticulation that we investigate. (A-B) Two hypothetical sequences of cursor movements, from the yellow to the blue and then the red nodes, without coarticulation (A) and with coarticulation (B). In the latter case, rather than moving directly from the yellow to the blue node, a shortcut to the red node is taken (note however that in our experiment, the path has to pass sufficiently close to the blue node to be considered valid). Hence, the kinematics of the first cursor movement are influenced by the next cursor direction (to the **East**) and carry some predictive information about it. (C-D) Two hypothetical sequences of saccade fixations, indicated by numbers, without cross-effector (gaze-cursor) coarticulation (C) and with cross-effector (gaze-cursor) coarticulation (D). In the former case, the fixation points are close to the target nodes, without a specific direction. Rather, in the latter case, fixation points slightly deviate towards the next target. (E-F) Two hypothetical sequences of saccade fixations, indicated by numbers, without (E) and with gaze coarticulation (F). While in the first case the angular position of the first and the second fixation points (relative to the first target node) are different, in the latter case they are very similar.

## Results

### The problem solving task

The participants solved 90 problem solving tasks, each requiring them to find a path that connected all the “gems” on a grid, without revisiting any nodes (though they were allowed to backtrack to unselect nodes) (Figure 1). Each problem had to be completed in 60 seconds, and more points were awarded for faster solutions. The participants solved the problems by controlling a computer mouse, while the grid and the cursor were always visible on a PC screen.

### Participants segment problems into sequences of gestures and alternate gaze and cursor movements

Our first question regards the possibility that sequences of gaze and cursor movements are organized in time, e.g., they alternate. Various previous experiments reported gaze anticipatory behaviour – or in other words, that gaze precedes the hand (Land, 2004; Land et al., 1999). However, despite being frequently observed, this anticipatory behavior is not general and depends on various conditions, such as the uncertainty of the task (Ambrosini et al., 2011). This requires assessing whether an anticipatory gaze regime is present in our task.

Our analysis reveals that the solution of our planning problem often comprises two phases, see Figure 3A for an example. In the *(pre)planning* phase, which is not included in the analysis, gaze (in blue) scans the problem while the cursor (in orange) stays still. In the subsequent *execution* phase, which is the object of our current analysis, gaze-cursor coordination follows an alternating pattern: gaze position anticipates the cursor position most of the time and tends to remain fixed on (or close to) a target node, until the cursor reaches it (or the cursor-fixation distance becomes very small). The alternating pattern can be visually appreciated in Figure 3D, which shows that cursor-fixation distance increases (as the gaze moves from one target to the next, while the cursor tends to remain stationary) and then decreases (as the cursor moves towards the next target node, and the gaze tends to remain stationary), multiple times.

**Figure 3:**
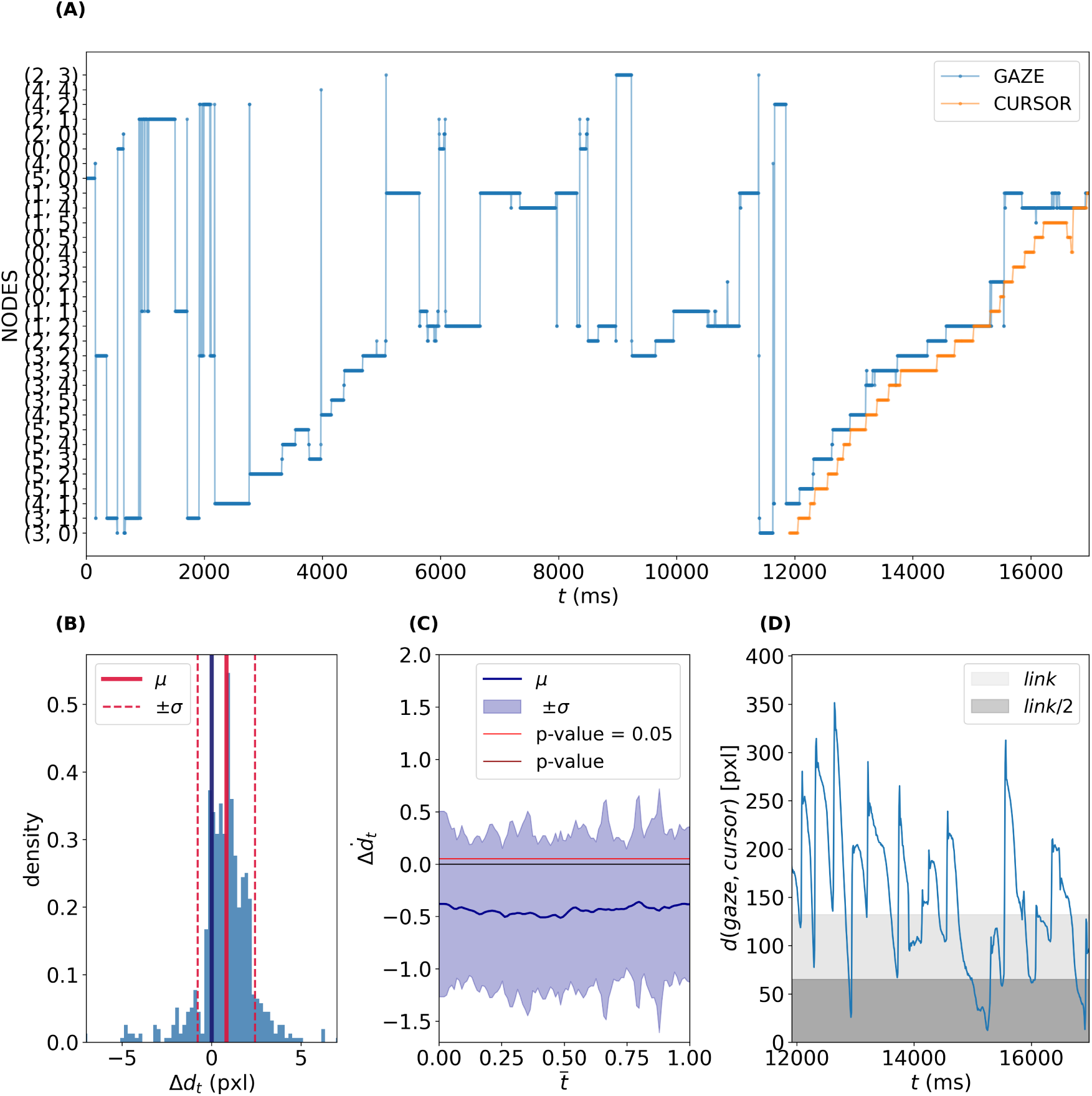
Gaze fixations and cursor movements during the solution of an example problem. (A) A visualization of the gaze-cursor coordination during the solution of a problem. On the y-axis nodes of the graph named after their coordinates; on the x-axis time. The blue and orange lines represent gaze and cursor positions collapsed onto the closest node of the graph. Note that our analysis focuses on the *execution* phase, in which participants execute both gaze and cursor movements – and which in this example starts around 12000ms. (B) Difference in gaze-cursor euclidean distance between consecutive fixations. (solid red line) *µ* is the mean of the distribution; (dashed red line) *σ* the standard deviation. (C) Temporal derivative of the distance between the gaze and the cursor position during a fixation over a normalized time (percentile of the fixation duration). (solid blue line) *µ* is the mean of the distribution; (shaded blue area) *σ* is the standard deviation interval; the (darker) red line is the estimated *p_value_* at each time (one-sample t-test), and the (lighter) red line is a threshold *p_value_*(*t*) = 0.05. (D) gaze-cursor euclidean distance during the implementation phase; *link* and *link/2* represent the full and half distance between nodes.

To test the statistical significance of a regime that alternates gaze fixations and cursor movements to the same target, we considered the variation in gaze-cursor distances between two consecutive fixations, defined as:

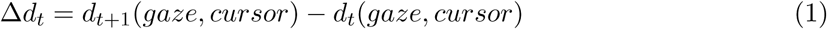

Positive (negative) values of this difference imply that fixations move away from (towards) the current cursor position.

The variation in gaze-cursor distances between consecutive fixations was on average significantly greater than zero (*τ* = 14.21*, p_value_ <* 10*^−^*^3^, one-sample t-test), implying that fixations move away from the current cursor position (Figure 3B).

We next tested whether gaze-cursor distance Δ*d* decreases during a fixation, as expected if gaze precedes the cursor on the same target. We computed the temporal derivative of Δ*d* during a fixation: a negative (positive) value in the derivative implies that gaze-cursor distance decreases (increases) over time. To compare distances during fixations having different duration, we first defined a normalized time *t̅* given by the absolute time *t* shifted to start from 0 and normalized by the fixation duration:

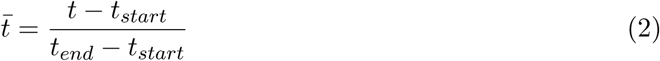

where *t_start_* and *t_end_* are the start and end of the fixation, respectively. The normalize time *t̅* indicates the percentage of fixation duration and is comprised between 0 and 1.

The time derivative of gaze-cursor distance was negative throughout the fixation duration (*p_value_*(*t̅*) *<* 10*^−^*^3^, ∀*t̅* ∈ [0, 1]), implying that, on average, the cursor reduces its distance from gaze position, therefore reaching the gaze on the same target (Figure 3D).

To summarize, during movement execution, gaze and cursor show alternating dynamics, with gaze fixating a target node until the cursor reached it – and then fixating the next target node. This alternating structure segments the problem into a sequence of *gestures*, defined as the intervals of time between two consecutive fixations, during which gaze remains stable on the target node while the cursor reaches it. The *gestures* identified in this analysis provide an organizing principle for the next analyses in the subsequent sections.

### Coarticulation in sequential cursor movements: both gaze position and current cursor kinematics predict the next cursor direction

We next aimed to assess whether participants coarticulated two subsequent cursor movements. Coarticulation generally involves executing the first action in a plan in a way that depends on the subsequent action in the plan. To assess coarticulation, we asked whether gaze position—while fixating on the current target—and the kinematics of cursor movement leading up to the current target contained information about future actions, such as the next direction of cursor movement (e.g., left, right, or straight after reaching the current target), and allowed to *predict* it.

To cast the analysis as a classification problem, we collapsed cursor trajectories after the current node into four categorical directions: **N**orth, **E**ast, **W**est or **S**outh. Then, in order to compare gestures of different length, duration, starting and ending nodes, we performed a standardization of their kinematics by fitting gestures with splines, resampling a fixed number of points along the spline, shifting it such that the starting node is placed in (0,0) and rotating it such that the first link to be crossed is the **N**orth. Even though participants were allowed to backtrack during the task execution, we decided for simplicity to exclude backtracking phases from the analysis. Thus **S**outh is not an available direction. Next, we used a method called *spatio-temporal Principal Components Analysis* (st-PCA) to obtain lower-dimensional description of the trajectories. This method has already been succesfully used in motor-control (Klein Breteler et al., 2007; Russo et al., 2014; Maselli et al., 2017, 2025) as decomposes the kinematics in simple and interpretable way. See Materials and Methods for the details and Supplementary Materials for the interpretation of the PCs.

Since the analysis above reveals that problem solving is naturally decomposed into a sequence of *gestures*, we measured prediction performance separately for the cases *within gestures* and *between gestures*. Predicting cursor direction *within gestures* implies that participants coarticulate consecutive movements to their current target, whereas predicting cursor movement *between gestures* implies that they coarticulate movements to the next target while they reach their current target. This procedure results in two predicting variables (gaze/kinematics) in two conditions (within/between gestures: four prediction tasks overall. Since we collapsed the next cursor direction into discrete values, we were able to solve these four tasks using standard classification methods. In this study, we used *Linear Discriminant Analysis* (LDA), a well-known linear classifier that has been previously applied to kinematics analysis (Maselli et al., 2017) with the same objective. Our goal is to evaluate how well an LDA model classifies gaze position and cursor kinematics into the three possible next directions (Figure 4A).

**Figure 4:**
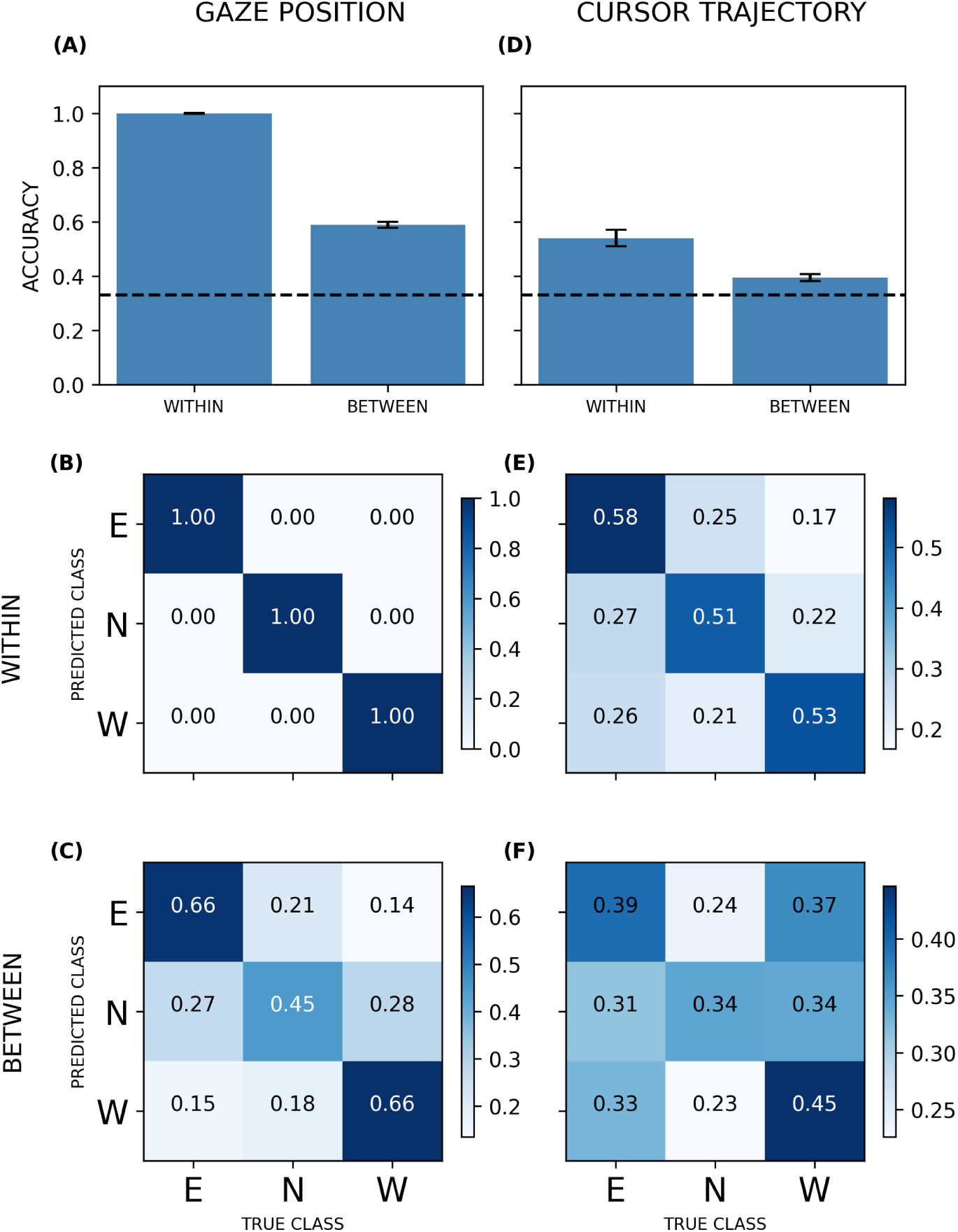
Quantification of the coarticulation effect in the next cursor direction. (A, D) Accuracy of the LDA models *within* and *between* gestures for (A) the gaze position and (D) the cursor trajectory. Error bars are the standard deviation obtained across 10 replicas of the crossvalidation. (B-C; E-F) Confusion matrices for the LDA models *within* and *between* gestures for all classes (*E*, *N*, *W*) for (B-C) gaze position and (E-F) Cursor trajectory.

We found that both gaze position and cursor kinematics predict the next cursor direction above chance (dotted line), both *within gestures* and *between gestures*. Two control analyses confirm these results: one shows consistency across all three game levels (Figure S4 and Figure S3), and the other includes gestures where the cursor’s endpoint does not match the gaze target—such as cases where the cursor left the current target before reaching it (Figure S6)—which were excluded from Figure 4A.

To obtain more information about the quality of the classification, we plotted confusion matrices for the LDA models corresponding to the four prediction tasks (Figure 4; see also Figure S7 for the control analysis. Confusion matrices entries show how many times a direction was correctly predicted or wrongly assigned to another value. The more the values of each row are concentrated in the diagonal, the better the discriminability of the directions. In all cases except one (i.e., cursor kinematics in the north direction between gestures) the diagonal shows greater values than the off-diagonal terms, implying a discriminability above chance.

To summarize, we found coarticulation in cursor-cursor and gaze-cursor movements, when participants made consecutive movements *within gestures* and *between gestures*. These results imply that participants plan ahead multiple cursor movements to both the current and the next targets.

### Coarticulation in sequential gaze fixations: the current gaze angular position predicts the next saccade direction

We finally asked whether the angle formed by the gaze position and the last node of a gesture shared information with the next fixation angle – which would imply the coarticulation between sequential fixations (see Figure 5 for the definitions of the variables). In other words, we tested wether there was a change in the relative position of the gaze with respect to the target node, depending on the direction of the next saccade (Figure 2E-F). Using the last node of a gesture as a reference for this analysis is a natural choice, as it is the point that defines the segmentation of sequential gestures.

**Figure 5:**
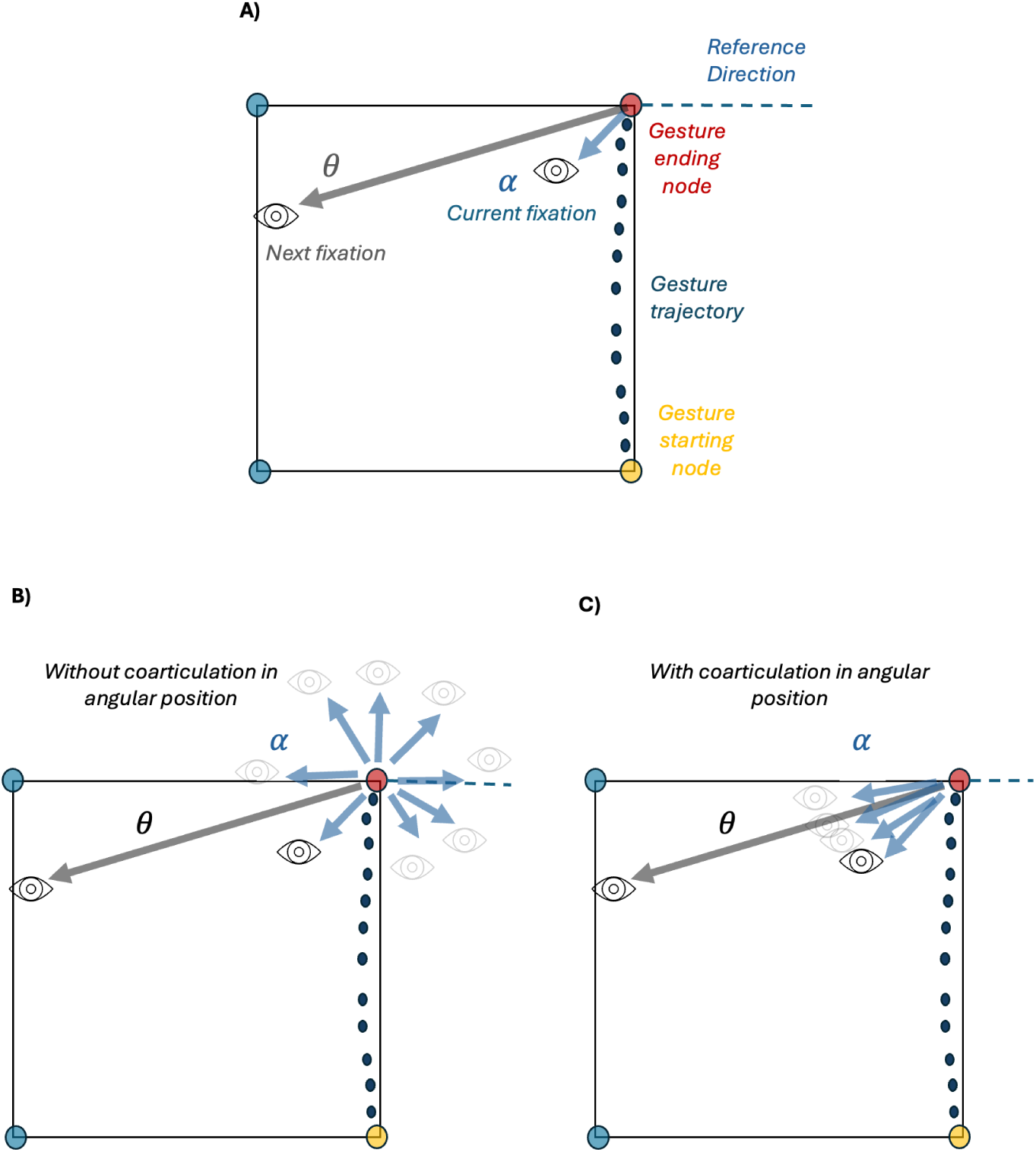
Definition of the frame of reference for the angular variables used in the analysis of coarticulation in gaze. A) A synthetic example of gesture connecting two nodes: (yellow) is the starting node of a trajectory (dark blue dots) that reaches a (red) ending node. The eye symbols indicates the gaze position: the (light blue) vector indicates the direction connecting the ending node to the current fixated position, thus identifying an angle (*α*) between this vector and the reference direction; analogously the gray vector connects the ending node to the next fixated position and identifies an angle (*θ*) with the reference direction. (B) A synthetic example of a sequence of fixation where there is no correlation between the current fixation angle and the next one. (C) A synthetic example of a sequence of fixations where the current fixation angle and the next one are correlated.

The pipeline used to study the gaze-to-gaze coarticulation is illustrated in Figure 6. Since we are fitting a relation between two angular variables-the angle formed with the target node of the current gesture by the current fixation and the next one-we resolved to use a so-called circular-to-circular model (Jammalamadaka and SenGupta, 2001) to regress current fixation angle to the next fixation angle (Figure 7A) as opposed to the most common linear-to-linear regression. Since there is no obvious goodness of fit measure for this circular-to-circular regression, we tested this model on a classification task. For this, we discretized the regressed output of the model and the true next fixation direction into 8 bins and then estimated the classification accuracy of this model. We included two controls in our analysis (Figure 7(B)): a model which regresses random points around the remaining (unvisited) nodes to the current fixation angle and a model regressing random points in an interval around the remaining (uncollected) rewards to the current fixation angle. The two control models only use available visual information (i.e., node and gem positions). The accuracy of the three models is greater than chance (dotted line; 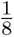), but the regression to the next fixation angle shows significantly greater prediction accuracy compared to the two control models (*A_Next_* = 0.350 ± 0.008*, p_value_ <* 10*^−^*^3^*A_Unvis_* = 0.183 ± 0.006*, p_value_ <* 10*^−^*^3^*, A_Uncoll_* = 0.225 ± 0.006*, p_value_ <* 10*^−^*^3^) (Figure 7B).

**Figure 6:**
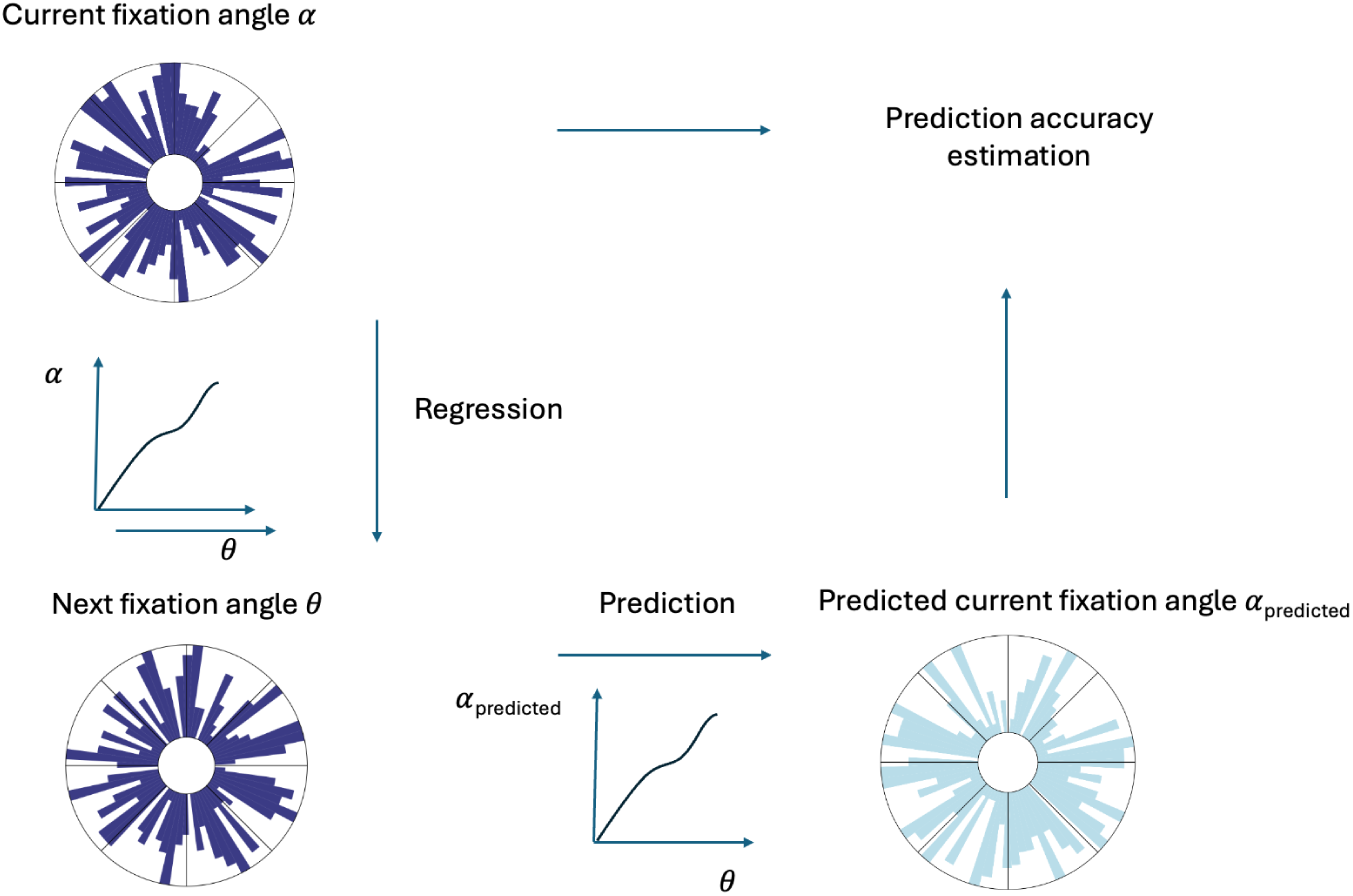
Gaze-to-gaze coarticulation analysis pipeline. This (synthetic example of) angle distributions are represented as (circular) histograms. The sectors in which the circles are divided correspond to the next binning into 8 discrete angle intervals, that we will use to test the model prediction accuracy. We first train a model to regress the current fixation angle to the next fixation angle. Given the next fixation angle as input, the model returns a prediction on what -according to this fit- current fixation angle would be. We finally discretize the angle values into the 8 bins to compare the prediction with the actual angle and estimate an accuracy of this. The control models follow the same pipeline but regress a different set of angles to next fixation angles (see main text for the details on these random model).

**Figure 7:**
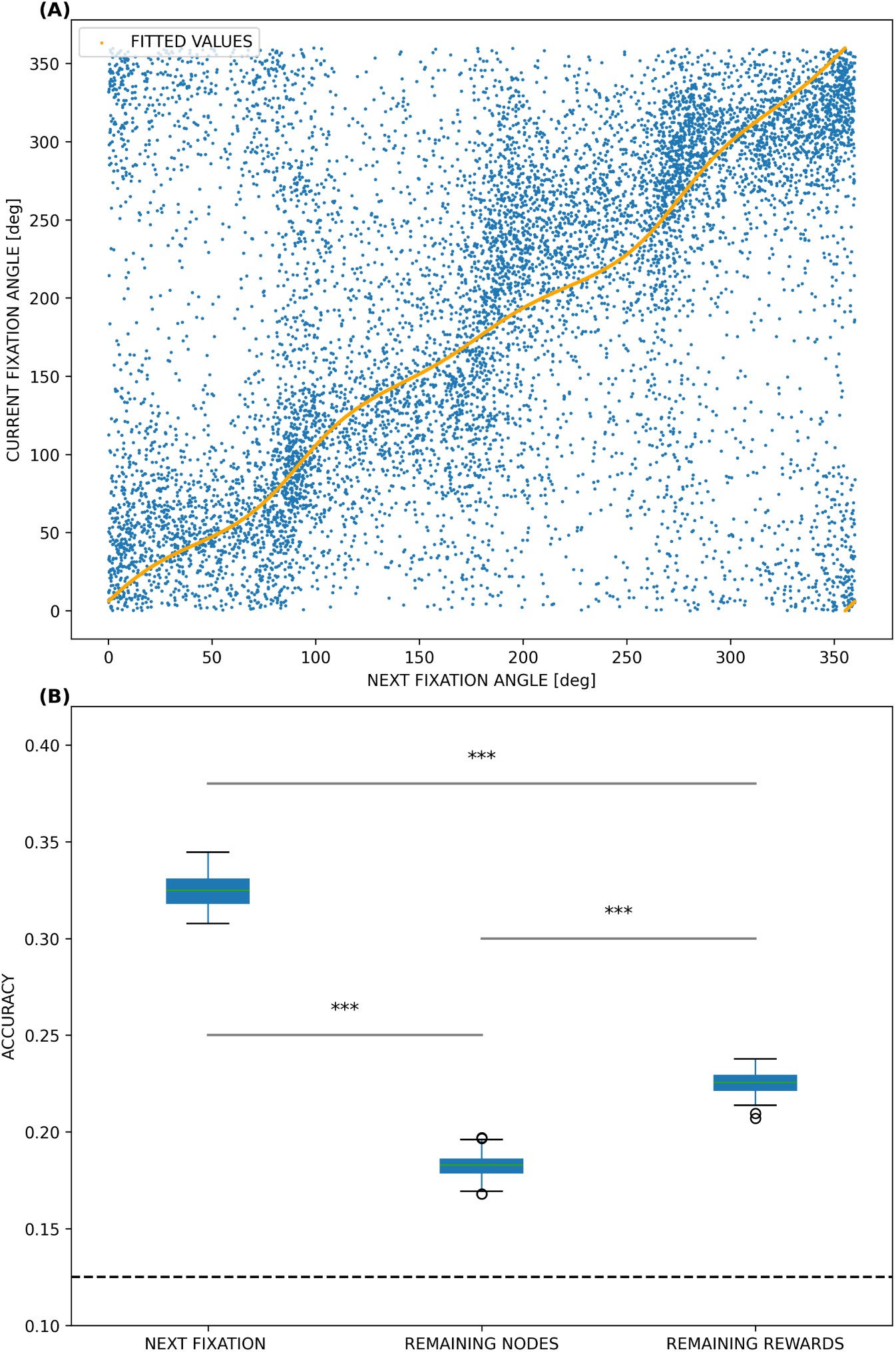
Quantification of the coarticulation effect in the next saccade direction. (A) Relation between the current fixation angle and the next fixation angle, in a Cartesian plane. The orange line represents the fitted values of the circular-to-circular regression. (B) Accuracy of the circular-to-circular regression model in the general case. The boxplots represent the accuracy of the models. The dashed line is the chance level.

To summarize, we found coarticulation between consecutive gaze fixations, indicating that participants plan ahead fixations to multiple targets.

## Discussion

Assessing how we plan sequences of actions to solve challenging problems has been a central question of cognitive science since its inception (Newell et al., 1972). Here, we investigated human planning strategies by recording cursor and gaze kinematics while participants solve problem solving tasks requiring navigating in a grid with the computer mouse, to collect multiple “gems”. We used the coarticulation of cursor-cursor, gaze-cursor and gaze-gaze movements as an index of planning. The rationale is that, to the extent that participants plan ahead multiple movements, the kinematics of their first movement in a sequence should carry information about – and hence predict – the next movements.

We report three main findings. First, after a (pre)planning phase in which the eyes scan the problem while the cursor remains fixed at the start position, gaze fixations and cursor movements show a robust alternating pattern, with gaze selecting a target node and fixating it until the cursor reached it – and then moving to the next target node. This finding suggests that plan execution is naturally segmented into a sequence of *gestures* – defined as the interval of time between two consecutive fixations.

Second, both gaze position and cursor kinematics predict the next cursor direction, which implies coarticulation both within a single effector (cursor-cursor) and across effectors (gaze-cursor). For example, the cursor movements directed towards the next node typically deviate from the beginning in the direction of the next node (Figure 2). Similarly, gaze fixations do not simply lie close to the next target but were also slightly deviated in the direction of the next target. Interestingly, not only coarticulation effects arise within gestures, but also between consecutive gestures. This finding indicates that participants plan ahead multiple cursor movements, to reach both the current and the next targets.

Third, and interestingly, the fixation angle with respect to a fixated node is consistently predicted by the next fixation angle, implying coarticulation in gaze. This novel finding indicates that while participants segment the task into a sequence of targets to be fixated and then reached with the cursor, they plan fixations to multiple targets in advance. Although sequential saccade planning has been previously reported (Zingale and Kowler, 1987; Hoppe and Rothkopf, 2019), here we show for the first time that the first fixation in a sequence is already positioned in the direction of the next fixation – hence predicts it. These results demonstrate that eye-hand coordination is not merely ‘the gaze guiding and anticipating’ hand movement; rather, future plans directly influence both the gaze and the hand movements. Furthermore, we show that the sequential gaze strategy is present during the solution of novel and challenging problems that are neither instructed nor routinized, but rather have to be found from scratch.

Overall, these results suggest that participants form hierarchically organized plans to coordinate eye and hand movements required for problem-solving. Participants segment the problem into a sequence of gestures and then achieve them by alternating between gaze and cursor movements. Specifically, gaze selects the target for each cursor movement (i.e., the endpoint of a gesture) and remains fixed on it to guide the cursor until the target is reached, then transitions to the next target. This alternation implies a hierarchical planning structure, with a higher level organizing sequences of gestures to define targets (or subgoals) for gaze and cursor, while a lower level supports the detailed sequence of cursor movements needed to reach each target. The hierarchical structure also reflects a separation of timescales, with gaze movements occurring at much slower intervals, serving as a framework within which the rapid pace of cursor movements is nested (Botvinick et al., 2009; Pezzulo et al., 2018). Notably, the coarticulation analyses indicate that participants not only plan multiple movements within a gesture but also plan multiple gestures in advance. This finding suggests that sequential planning operates simultaneously at both the higher (gesture) and lower (cursor movement) hierarchical levels, supporting complex coordination across the task.

One limitation of the current study is that – since we were interested in the structure of the coordinated eye-cursor plans – it only focuses on the action *execution* phase. Future studies could address the relations between the (pre)planning phase, in which eye movements scan the problem to establish a (partial) plan and the subsequent execution phase. Furthermore, future studies could investigate the events that break the alternation of gaze and cursor movements during the execution phase (e.g., pauses, backtracking, movement errors), which we excluded from the analysis. These events indicate that planning does not end up when movement execution starts, but rather continues afterwards, in parallel or in alternation with movement execution (Nuzzi et al., 2024). This explains also why we called the phase occurring before movement *(pre)planning*. Future studies of the events that break the alternation of gaze and cursor movements could reveal more fine-grained aspects of planning and replanning during problem solving. Finally, future studies might use our problem solving paradigm to advance our understanding of the neural underpinnings of action sequencing and hierarchical planning in humans and other animals (Crivelli-Decker et al., 2023; Saito et al., 2005; Dehaene et al., 2015; Geddes et al., 2018; Balaguer et al., 2016).

## Materials and Methods

### Experimental Setup and Procedure

We recruited 31 participants (19 M age = 27 ± 4 years, 12 F age=24 ± 5 years) with normal-to-corrected vision. Participants were free to leave the experiment at any moment. All gave informed consent to our procedures which were approved by the Ethics Committee of the National Research Council. All of them were rewarded for taking part to the experiment with vouchers. All the participants completed the experiment.

Participants were shown a video tutorial and informed about the task structure (4 training problems followed by 90 problems divided into 3 levels of 30 problems each (Eluchans et al., 2024)).

We used an Eye-Link 1000+ tracker in a chin-rest desktop mount configuration with a 35 mm lens. The experiment was performed inside an isolated room with no source of light except the IR camera of the gaze tracker and the (LED) light of the screen where the task app was running.

Participants could control the movement in the PC via a mouse cursor.

For each participant, a standard 9+1 points calibration was performed on a grid with the same area of the task problems at the beginning of the experiment. A 9+1 point validation was performed after each calibration. Participants were asked to move as little as possible once the calibration was validated. The acceptance criterion for the calibration was the one suggested by the producer: a maximum error in calibration of 1° and an average error smaller than 0.5°. Eye-tracking starting sampling frequency was 2000 Hz, while mouse-tracker sampling frequency was 60 Hz.

### Data Preprocessing: cursor trajectory resampling and dimensionality reduction

To enable comparison across gestures of varying durations (and because of the fixed sampling frequency, of varying number of points), we fitted a linear spline with no smoothing factor to each original trajectory and resampled a fixed number (N = 50) of 2D coordinates. Resampled points were spatially distributed according to the original velocity profile, ensuring that regions with fewer points (due to higher speeds) maintained the same relative density in the resampled trajectory.

The resampled trajectories were then translated such that the starting node coincided with the the origin of a Cartesian plane (0,0), and rotated such that the first crossed link direction was along the positive y-axis (“North” direction). The gaze position during the execution of this trajectory was described by a 2D point in the Cartesian plane and transformed (translated and rotated) according to the same amounts of the corresponding trajectory.

After standardizing each trajectory into a 2(Cartesian) x50(Sampled points) dimensions, we looked for a lower-dimensional description via a *spatio-temporal principal component analysis* (st-PCA). The st-PCA is a method that explains trajectory variability by identifying a basis of trajectories, or *spatio-temporal principal components*, whose linear combinations can reconstruct the original trajectories.

At the base of this method is the idea to treat each point of the (resampled) trajectory as a distinct variable. The 2D coordinates are then flattened in a single vector, resulting in each trajectory of N points being represented as a 2N-vector (*x*_1_*, …, x_n_, y*_1_*, …, y_n_*). Stacking all the considered trajectories creates an *N_trajectories_* × 2*N* matrix, where a standard PCA can be carried on.

Interestingly, we found the PCs emerged from the analysis of cursor movements to be functionally interpretable (Figure S9), supporting its appropriateness.

### Classification of the next cursor direction

We train four linear classifiers to predict the next cursor direction based upon two different effectors (the low-dimensional representation of cursor kinematics and gaze position) and for two different conditions (*within gestures* and *between gestures*).

For all the classifiers, we collapse the cursor directions into the three discrete orientations (**N**orth, **E**ast, **W**est) relative to the first movement, depending on the underlying sequence of nodes that are crossed (e.g. from (0,1) to (0,2) would correspond to the North direction).

The former, *within gestures* condition, considers whether the cursor kinematics up until a node can be used to predict the cursor direction towards the next node, when the two nodes belong to the same gesture. For the *within gestures* prediction, we consider all periods of time in which at least two links have been crossed and we use the cursor trajectory from start to the first node to train a classifier and the cursor trajectory from the first to the second node as the test for classification. The second, *between gestures* condition, considers whether the cursor kinematics up until a node can be used to predict the cursor direction towards the next node, when the two nodes belong to two consecutive gestures. For the *between gestures* prediction, we consider all the cases in which a single link was crossed during the first gesture (used to train the classifier) and at least one link was crossed during the second gesture. We exclude from analyses all the cases in which the participant paused or backtracked.

For classification, we use *Linear Discriminant Analysis* (LDA) that identifies a linear combination of features that optimally differentiate between two or more classes, while minimizing the variance within each class. To assess the statistical significance of the LDA model accuracy in the different cases, we used a *repeated stratified K-fold cross-validation* procedure (number of folds = 3, number of replicas = 100). For all the cases, we balanced the classes’ volume by a downsampling procedure, in order to avoid possible biases in the classifier. Since the downsample is a stochastic procedure we considered 30 (down)samples of the whole data, and for each of them performed the cross-validation procedure previously described. The errors estimated for the average accuracy are obtained as the standard deviation across all the (30 samples × 100 K-fold replicas) accuracies. This is a more conservative estimation with respect to taking the standard deviations of the (30 samples) accuracies.

### Gaze-to-gaze prediction

Finally, we study the information that the gaze position shares across consecutive fixations. In particular, during each gesture, we can define the current angular gaze position as the direction formed by the last node reached during that gesture and the current gaze position. Analogously we can define the next angular gaze position as the direction formed by last node reached during that gesture and the gaze position of the next fixation (Figure 5).

To fit the relation between these two angles we used a circular-to-circular regression (Jammalamadaka and SenGupta, 2001). This method fits separately sine and cosine of the dependent angular variable, with trigonometric polynomials of degree *m* and comes with a natural test for the optimal degree of *m*. However there is no simple way to estimate the goodness of the fit. We thus decided to evaluate the goodness of the fit by testing the accuracy of the classification of the dependent angle once discretized into 8 possible angular bins (Figure 6). We regressed the current fixation angle to the next fixation angle. Then we discretized the current fixation angle and compared the predictions of the regression model with the actual current fixation angle. Differently from the gaze-to-cursor case, in the estimation of this accuracy it was not used a downsampling procedure because classes were (approximately) balanced: the null case is 0.126, against 0.125 of a perfectly balanced case 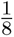. To obtain a more reliable estimation, the accuracy of the model we used a 3-fold cross-validation procedure repeated 50 times.

The estimated accuracy was then compared to two different random models. The former regresses the current fixation angle to the direction of a point randomly sampled around the nodes that had not been visited (rather then the next fixation position). The latter regresses the current fixation angle to the direction of a point randomly sampled in an interval around the “gems” that have not yet been collected. For the control models, we additionally considered 30 sets of randomly sampled variables (unvisited node for the first model, uncollected gems for the second one), and for each of them we performed a 3-fold cross-validation procedure repeated 10 times. The accuracy of the model was then computed as the average of the accuracy of the 30 instances of the models.

## Supporting information

Example video

## Author Contributions

Conceptualization: ME, GP; Methodology: ME, GLL, AM; Validation: ME, GLL, AM, GP; Formal analysis: ME; Investigation: ME; Software: ME, AM; Data Curation: ME; Writing - Original Draft: ME, GP; Writing - Review & Editing: ME, GLL, AM, GP; Supervision: GP; Project administration: GP; Funding acquisition: GP.

## Author Declaration

We have no competing interests to declare.

## Acknowledgments

This research received funding from the European Research Council under the Grant Agreement No. 820213 (ThinkAhead), the Italian National Recovery and Resilience Plan (NRRP), M4C2, funded by the European Union – NextGenerationEU (Project IR0000011, CUP B51E22000150006, “EBRAINS-Italy”; Project PE0000013, “FAIR”; Project PE0000006, “MNESYS”), and the Ministry of University and Research, PRIN PNRR P20224FESY and PRIN 20229Z7M8N. The GEFORCE Quadro RTX6000 and Titan GPU cards used for this research were donated by the NVIDIA Corporation.

## Supplementary Information

### Additional details about the planning task

The planning task comprises 90 problems, organized into three interleaved blocks (referred to as “levels”), with each level containing 30 problems. Each problem grid of the planning task was uniquely generated, with an average edge density of 0.75 ± 0.2 (meaning that, on average, 25% of possible edges were removed). This density was determined during a pilot study to offer a good balance of planning solutions. Before the experiment, participants completed a brief practice session with four problems, whose results were not analyzed.

We manipulated the planning demands both between and within levels. Between levels, planning difficulty increased with progressively larger maps and more gems to collect, making the higher-level problems generally more difficult. Within each level, the planning demands were varied by dividing the level into three sublevels, each consisting of 10 problems. These sublevels included problems that could be solved with three different planning depths. Planning depth for a problem was determined through simulation using a synthetic agent. The agent employed a breadth-first search strategy to find the shortest path that collected *n* rewards (e.g., all paths that collected 2 rewards when *n* = 2). The agent would then remove this partial path from the graph and repeat the process until either a complete solution was found (collecting all the rewards) or no more rewards could be collected (for instance, if the agent trapped itself in a dead end). A problem was considered to require planning depth *n* if this was the smallest value of the parameter that allowed the agent to find a solution.

Since the accuracy for the four classifiers (trained upon gaze position and cursor trajectories in the *within* and *between* gestures conditions) show no significant differences across the levels of the factorial design of the experiment, we combined the in the main analysis. Separate results for the three levels are shown in Figure S3 and Figure S4.

### Additional details about the experimental setup

We used an EyeLink 1000 Plus for eye-tracking. The eye-tracker configuration used was the chin-forehead rest desktop-mount and realized following the indications of the producer: the chin-forehead support was kept at a fixed height such that the eyes of the subjects were approximately at ¾ of the screen height. Subjects seated on a chair with adjustable height. The IR-camera was placed at approximately 50 cm from the eye position of the subject, between the subject and the screen. Subjects could control the movement in the app via a mouse cursor. The synchronization of the game app and the eye trackers was performed via a Python script mainly relying on Pylink libraries (the Python library of the Eye Link) and the PyAutoGUI (a Python library for mouse and visual interactions).

### Additional details about the experimental procedure

Each participant was first shown the video tutorial (https://drive.google.com/file/d/1RDkY6B_-vM830H_2HgZK1tELSA5N9vJq/view) and given the information about the structure of the experiment: (1) calibration, (2) validation, (3) task divided in three levels of 30 problems each, preceded by 4 training problems. Participants were instructed that their total score depended on problem completion and its velocity: the faster the completion, the greater the number of points they earned. Participants were also instructed that the “gems” could randomly appear with two colors (blue or red) with the latter case yielding double points. The passage from the trial levels to the first level, as well as the passages from each level to the next, and the end of the whole experiment, were indicated by a message on the screen.

For each participant, a standard 9+1 points calibration was performed on a grid with the same area of the task problems at the beginning of the experiment. A 9+1 point validation was performed after each calibration. Participants were asked to move as little as possible once the calibration was validated. The acceptance criterion for the calibration was the one suggested by the producer: a maximum error in calibration of 1° and an average error smaller than 0.5°. Eye tracking sampling frequency was 2000 Hz, while mouse-tracker sampling frequency was 60 Hz.

The duration of the experiment varied across participants, ranging from 30 to 60 minutes.

### Fixation detection

Fixations show a great variability in terms of duration, and can vary depending on the task being performed. We decided to accept fixations with a minimum temporal duration of the 80 ms.

### Velocity threshold and pause detection

A preliminary analysis of the distribution of cursor velocity reveals that it is bimodal (Figure S11). One peak corresponds to greater speed, when participants move across nodes, while the other peak corresponds to smaller speed, when participants do not move the cursor or move it very slowly. The threshold used to detect the end of a movement is set at the minimum between the two peaks. This threshold was thus computed for each participant. A pause was identified if the cursor speed fell below the participant-specific threshold for at least 50 ms.

### Exclusion criteria for the analyses

The alternated coordination of gaze and cursor movements identifies a segmentation into *gestures* of an otherwise continuous dynamic, affording the analysis of information shared between consecutive gestures. To reduce the sources of noise, we considered various exclusion criteria. For both the cursor classification and the circular-to-circular regression analyses, we excluded all the gestures during which a pause or a backtrack was performed. In the *between* cases, we only considered cases in which this criterion holds for two consecutive gestures.

For the cursor classification analyses, we also considered additional exclusion criteria. First, we only included the gestures during which the last (target) node reached by the cursor trajectory matched the fixated node. We therefore excluded from the analysis the cases in which the eyes fixated another target before the cursor completed the reaching movement to the current target. However, the same qualitative results reported in the main classification analysis also hold when these cases are included (Figure S6 and Figure S7).

Second, to avoid classification biases, when considering the cursor trajectory between two nodes we removed the last part of the trajectory, after the (invisible) node collider was crossed. The collider had a radius of approximately one third of a link between two nodes.

Third, since the classification needs to compare trajectories having the same structure, in the *within gesture* case, we only considered gestures in which at least three nodes were crossed (starting node, intermediate node, ending node). The first part of the trajectory (starting node to intermediate node) was used to train the classifier into the second part (intermediate node to ending node). In the *between gesture* case, we only considered consecutive gestures where both the first and the second gesture comprised at least two nodes (starting node, ending node).

The number of cases considered for the main analyses of the *within gesture* condition were *N_samples_* = 1359, divided into North (*N*) = 934, West (*W*) = 229, East (*E*) = 196. The number of cases considered for the main analyses of the *between* condition were *N_samples_* = 4063, divided into North (*N*) = 1489, West (*W*) = 1438, East (*E*) = 1136. Since we excluded the backtracking cases and standardized the trajectories, the South (*S*) case is not present.

In the control analysis in which also included the cases in which the eyes fixated another target before the cursor completed (Figure S6 and Figure S7), the cases for the *within gesture* condition are *N_samples_* = 3044, divided into *N* = 2224, *W* = 443, *E* = 377. For the *between gesture* condition, they are *N_samples_* = 9299, divided into *N* = 3375, *W* = 3149, *E* = 2775.

### Functional interpretability of principal components

We sought to investigate whether the principal components (PCs) emerged from PCA analysis of cursor kinematics are functionally interpretable. For this, in Figure S8 we plotted the projection coefficients of the first 10 principal components in the original space (i.e., the 2D 50 points-resampled trajectories, with the first 50 points being the *x* coordinates and the remaining 50 points being the corresponding 50 *y* coordinates), by combining together the *within gesture* and *between gesture* conditions. Interestingly, we found that the principal components are interpretable. The first two components represent lower-level components of movement; namely, an offset with respect to the average, with the second component that seems to show a mild dilation/contraction effect along the y-axis. In these and the following components, it is also possible to observe a also a clear alternation in the relevance of the x and y coordinates in the projection coefficients, which segregate their importance into different components. The next principal components represent more complex aspects of movement. The third and fourth component represent a curvature in the trajectory where there is a crossing of the vertical axis. The fifth and sixth components introduce curves where the beginning and the ending of the trajectory bend towards a direction, but the middle part of the trajectory moves toward the opposite. The information encoded in the third and fourth and in the fifth and six components seem complementary: the former encodes the shortest path (and thus bend the trajectory towards the next target) and the latter encoded smoothers and longer paths. Finally, higher components show a higher complexity, with more S-shaped trajectories. A similar pattern is observed when considering principal components emerged from the separate *between gesture* (Figure S9(A)) and *within gesture* (Figure S9(B)) conditions.

### LDA coefficients

Figure S10 shows an example of LDA coefficients, where we projected the PCs over the LDA space, for both the *within gesture* and *between gesture* conditions. The values show how much each principal component discriminates the three labels considered in the classification tasks. As the number of the components increases, the unexplained variance decreases and the relative discriminative power increases.

The coefficients shown in Figure S10 are for the LDA models trained using all the available data. These models are obviously prone to overfitting and are not the ones used for the cross-validation, hence the corresponding values should be considered simply as illustrative examples.

It is possible to notice that the first two components, corresponding to global offset with respect to the mean (approximately the vertical axis), do not contribute to distinguish the classes. Interestingly, most of the higher components exhibit two possible behaviours: either they disambiguate the “turn vs straight” choice (i.e., E/W vs N direction) as in the fourth or fifth components, or the disambiguate the “left vs right” choice (i.e., E vs W direction) as in the fifth or seventh component, and the N direction is in an intermediate position between E and W. Clearly, however, some differences exist between the coefficient for the three labels and the two cases,

The methods described below aim to assess the three types of coarticulation described above, by assessing 1) whether it is possible to predict the direction of the next cursor movement based on the kinematics of the current cursor movement 2) and/or the current angular gaze position and 3) whether it is possible to predict the next angular gaze position based on the current angular gaze position.

---

**Figure S1:**
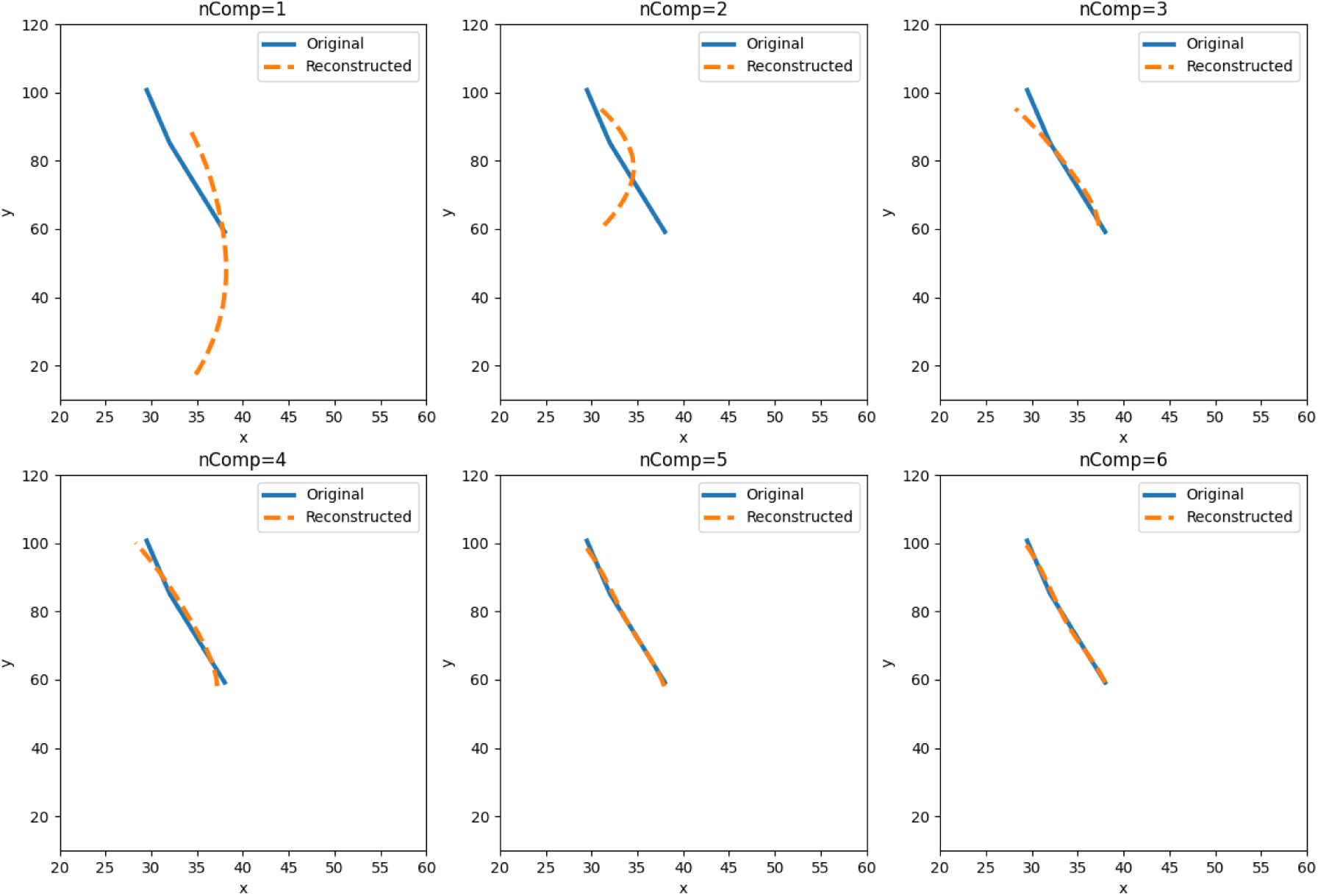
Reconstruction accuracy of cursor trajectories with PCA. An example cursor trajectory is reconstructed by progressively adding the first six principal components (PCs), with the original trajectory in blue and the reconstructed one in orange.

**Figure S2:**
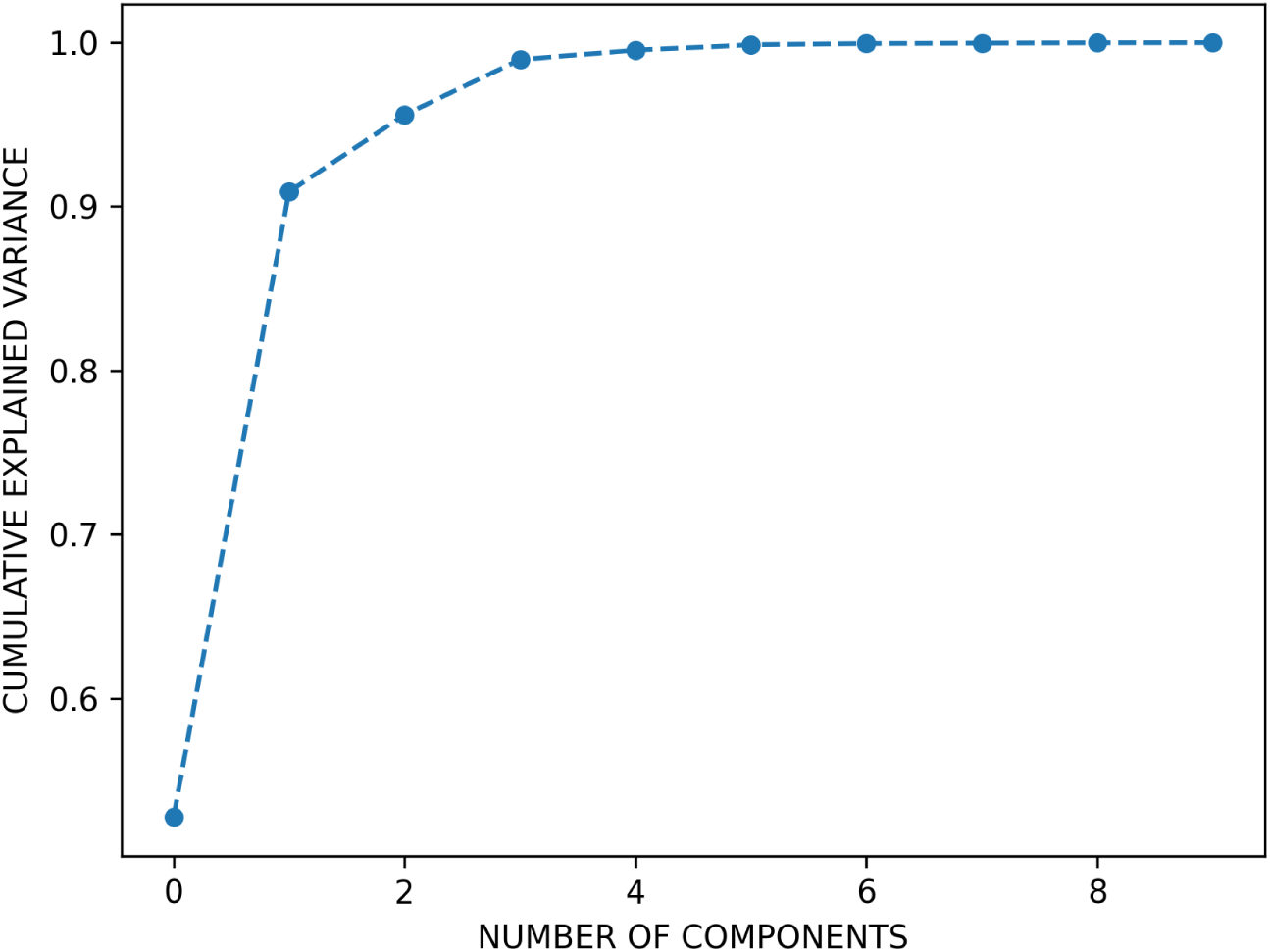
Reconstruction accuracy of cursor trajectories with PCA. Cumulative explained variance as a function of the number of PCs.

**Figure S3:**
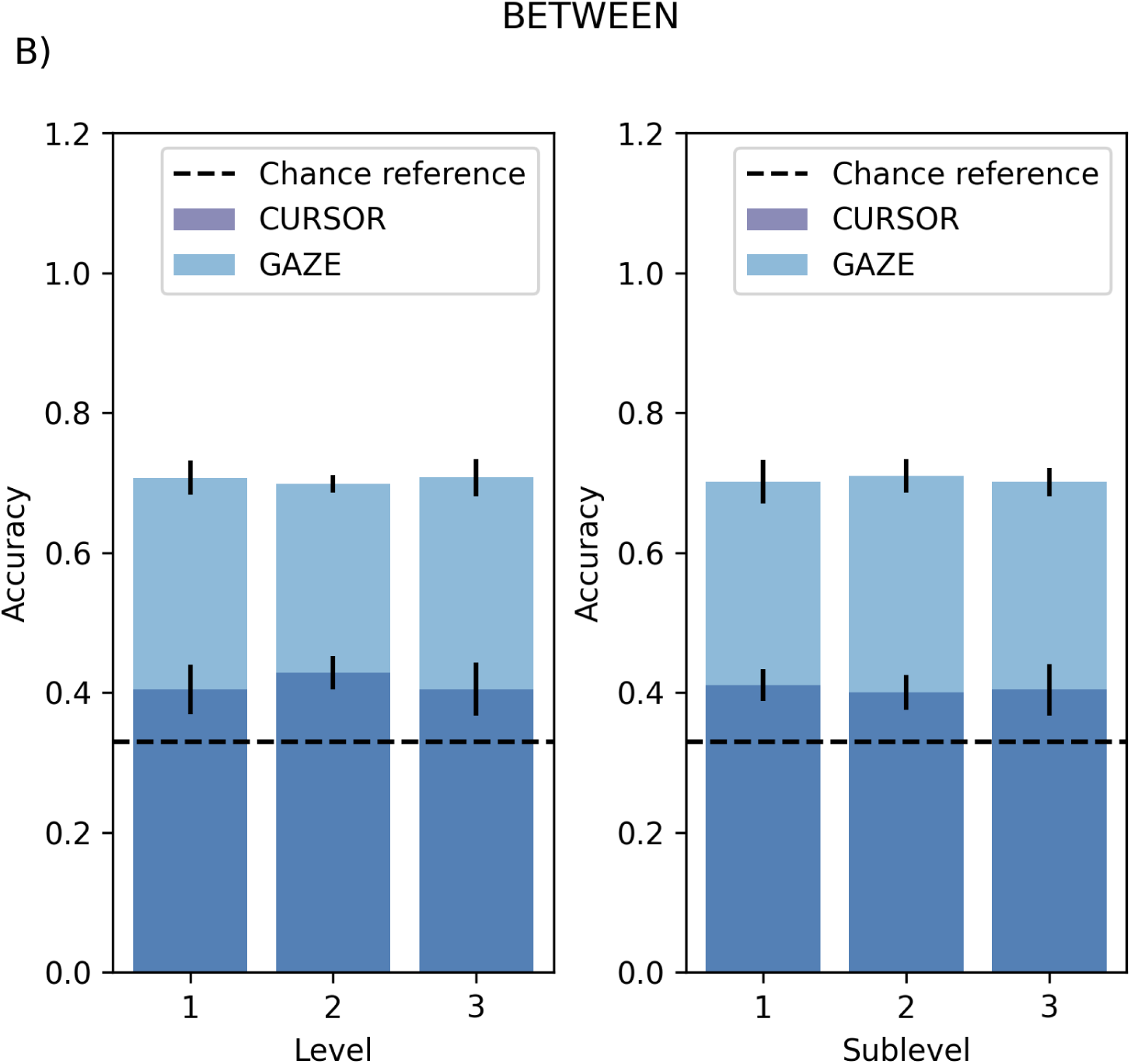
Coarticulation effect does not depend on the level/sublevel factors. Classification accuracy for gaze and cursor trajectories across the experiment factorial design (Level/Sublevel) in the *between* case.

**Figure S4:**
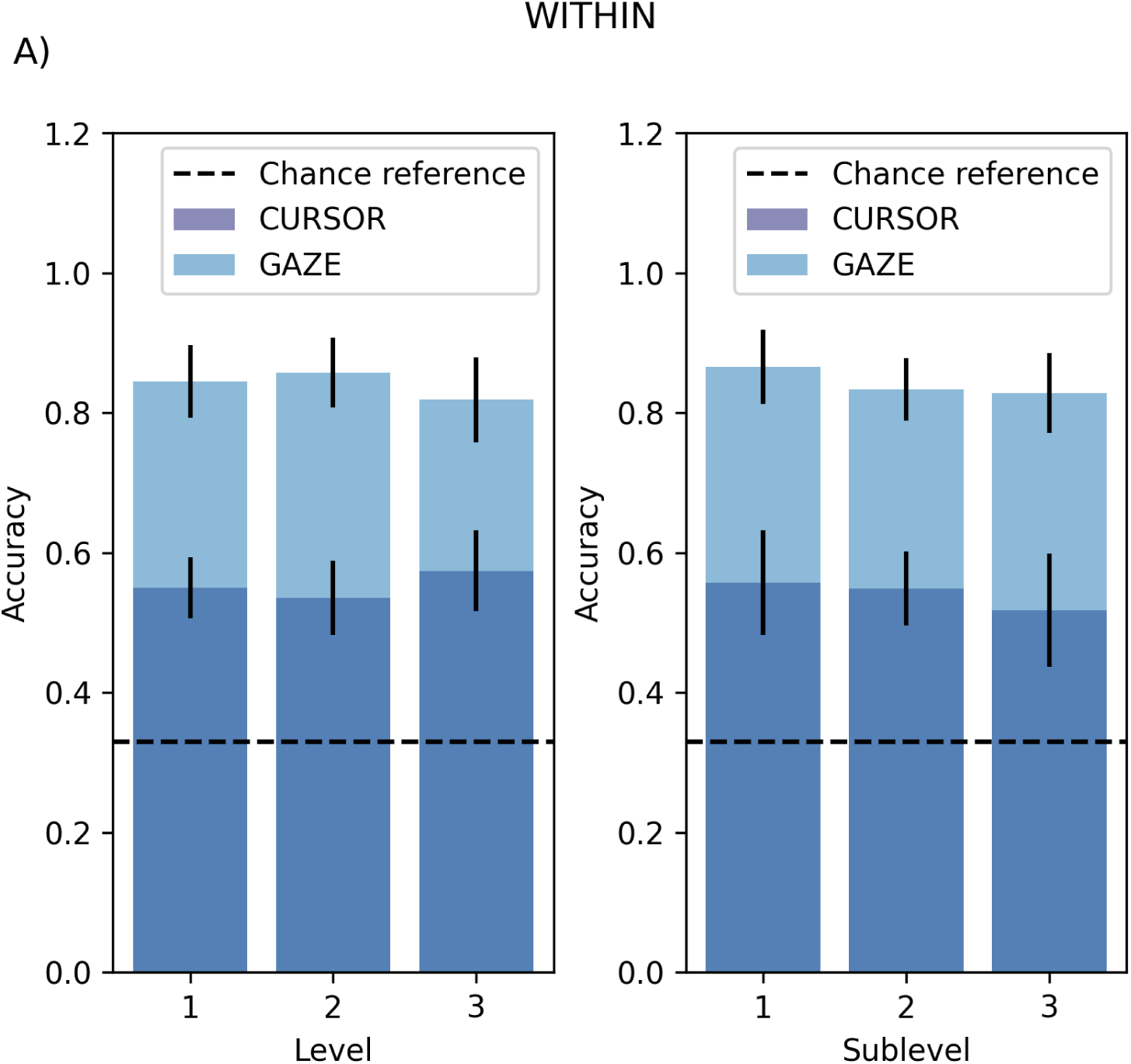
Coarticulation effect does not depend on the level/sublevel factors. Classification accuracy for gaze and cursor trajectories across the experiment factorial design (Level/Sublevel) in the *within* case.

**Figure S5:**
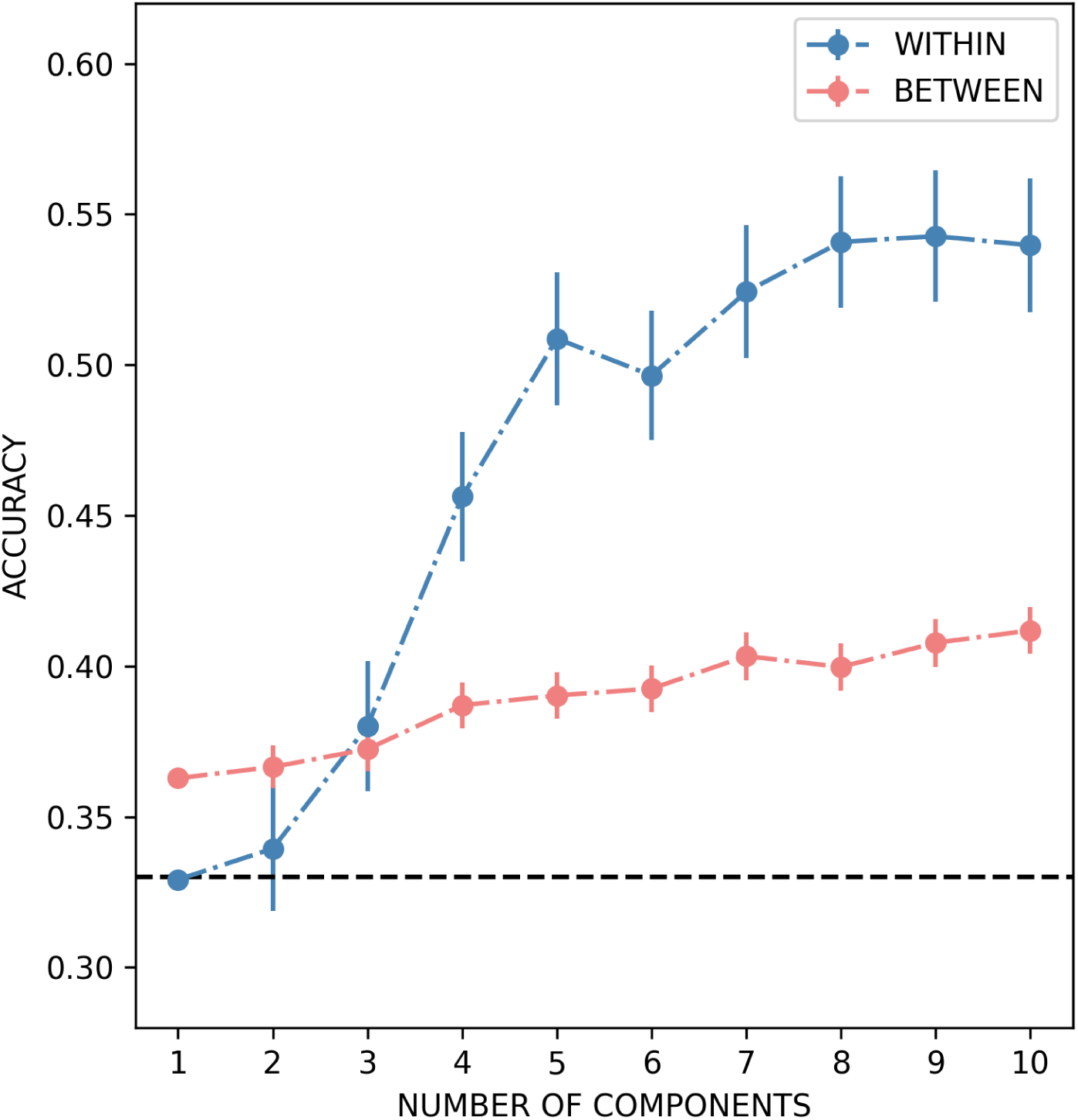
Quantification of coarticulation in cursor movements. The accuracy for LDA classifiers trained on trajectories compressed to a varying number of principal components (from 1 to 10) for the control case.

**Figure S6:**
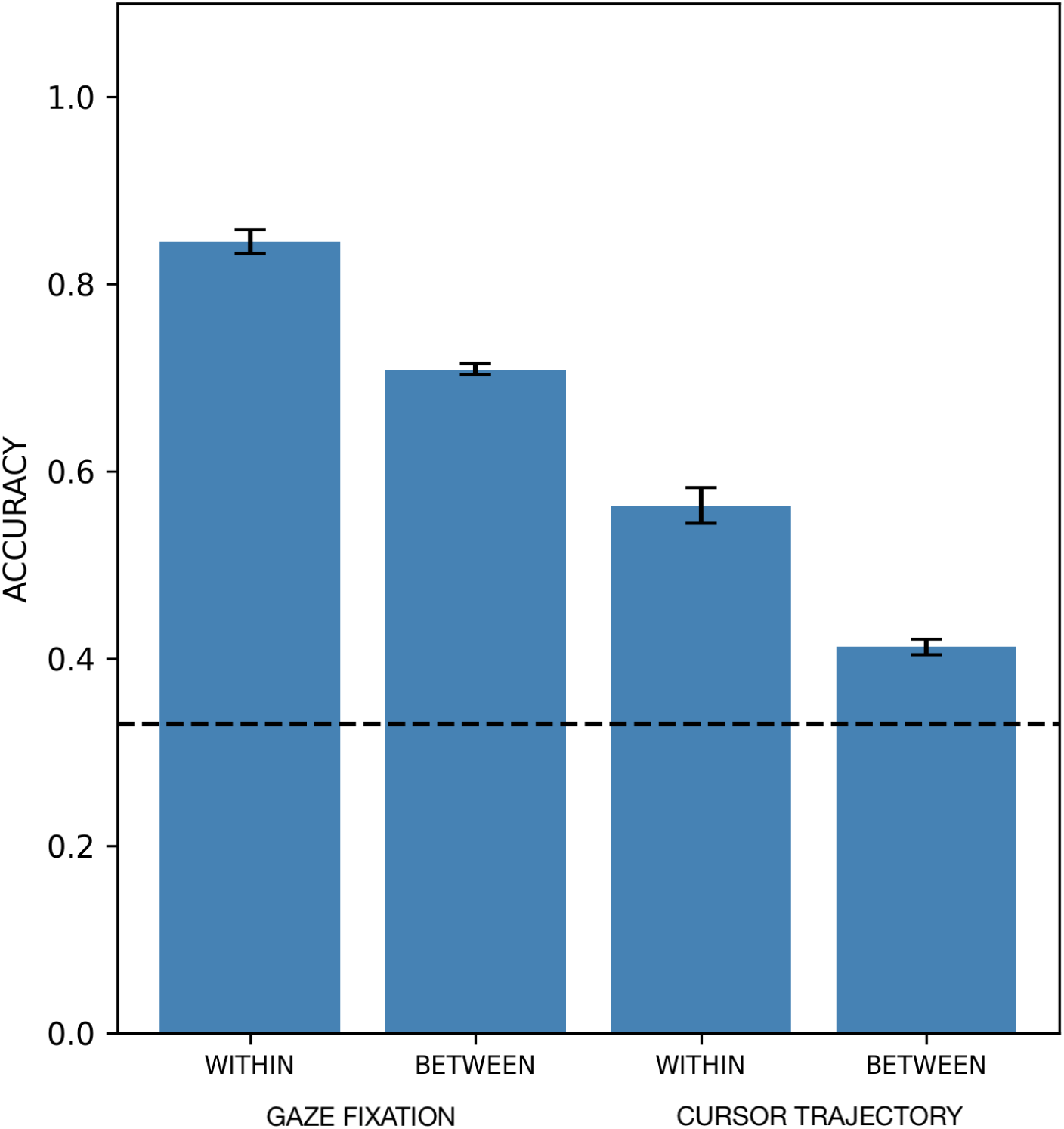
Quantification of coarticulation in cursor movements, in a control analysis that includes the cases in which the eyes fixated another target before the cursor completed (which are excluded in the main analysis). Accuracy of the LDA models in the different cases. The error bars represent the standard deviation of the accuracy across the 10 replicas of the 3-fold cross-validation.

**Figure S7:**
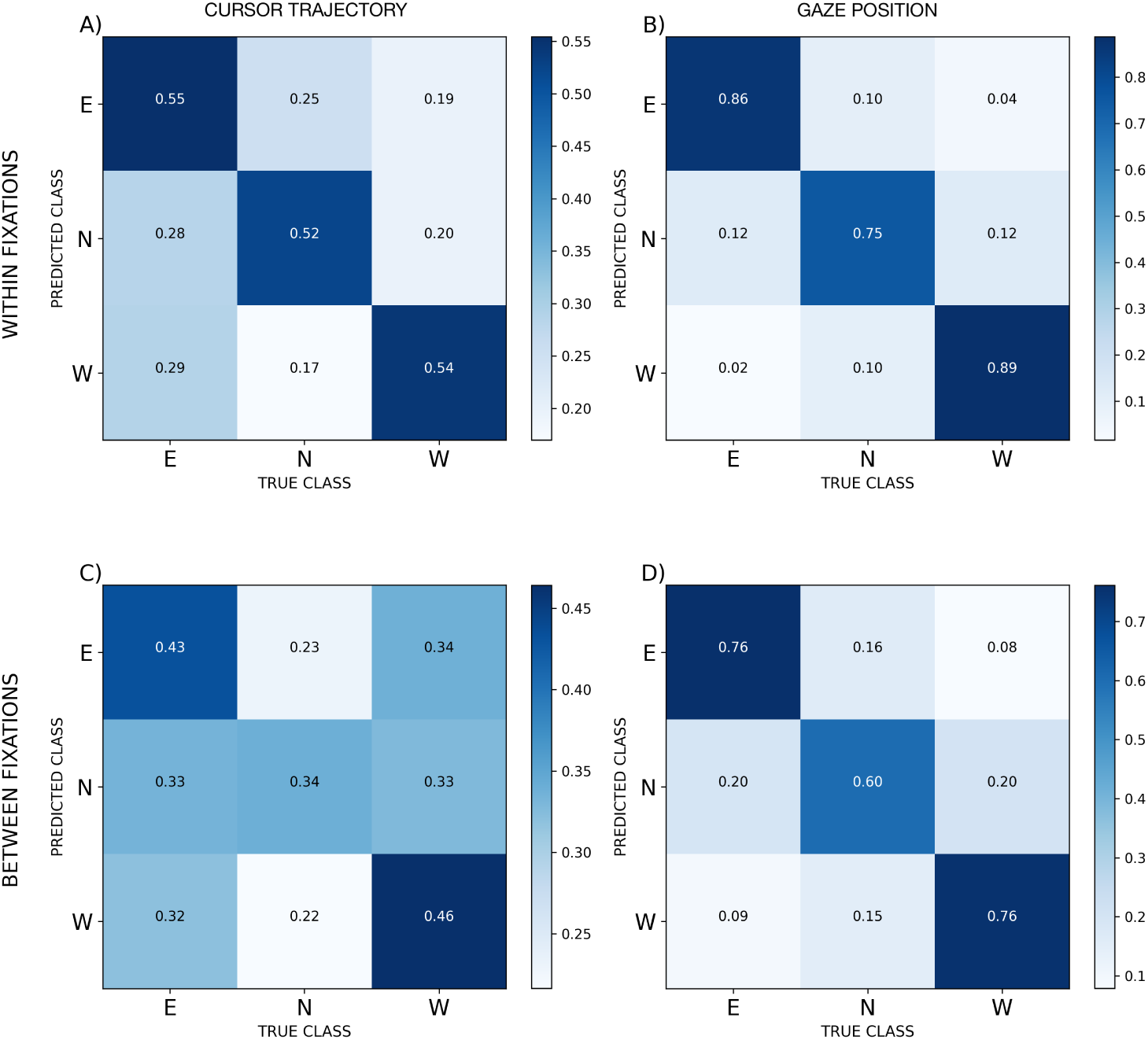
Confusion matrices for the LDA models in a control analysis that includes the cases in which the eyes fixated another target before the cursor completed (which are excluded in the main analysis). The values are the average of the confusion matrices across the 100 replicas of the 3-fold cross-validation.

**Figure S8:**
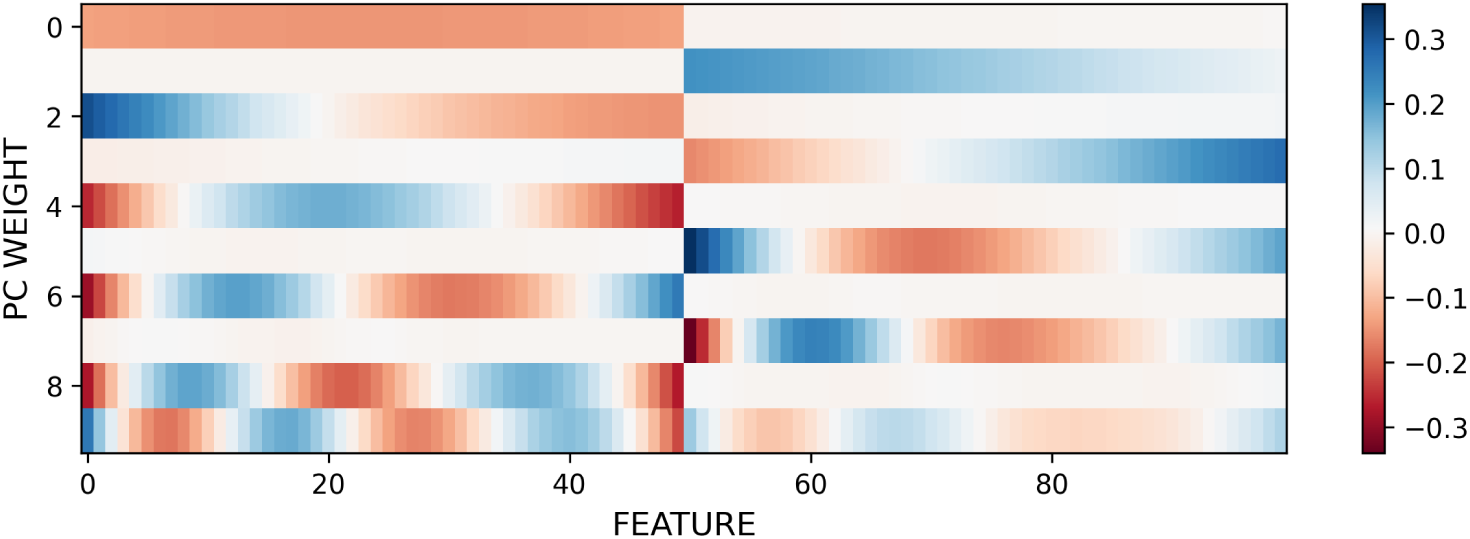
Spatio-temporal principal components interpretability. The panel shows the projection coefficients of the first 10 principal components in the original space.

**Figure S9:**
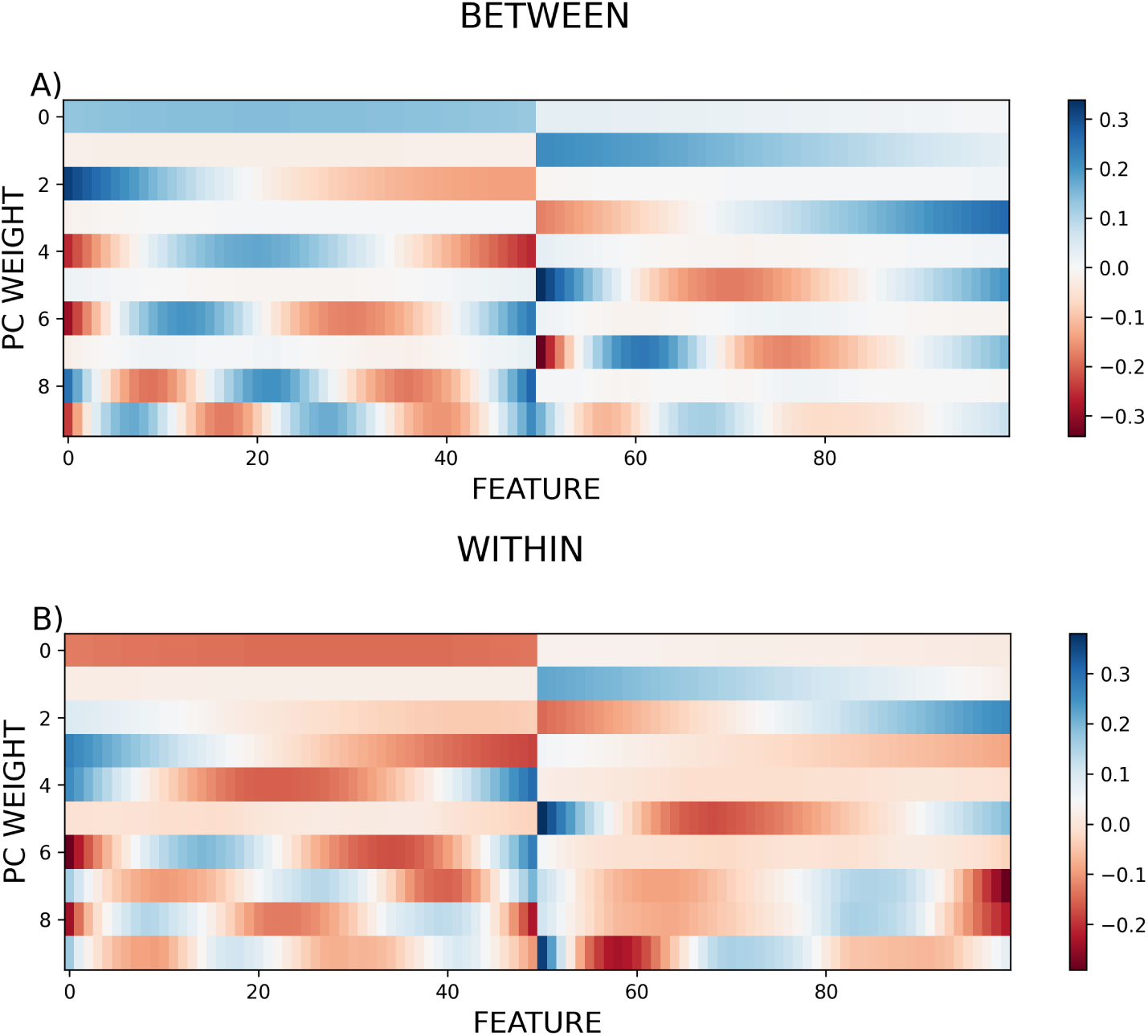
Spatio-temporal principal components interpretability. The panel shows the projection coefficients of the first 10 principal components in the original space for the (top) *between gesture* condition and the (bottom) *within gesture* condition.

**Figure S10:**
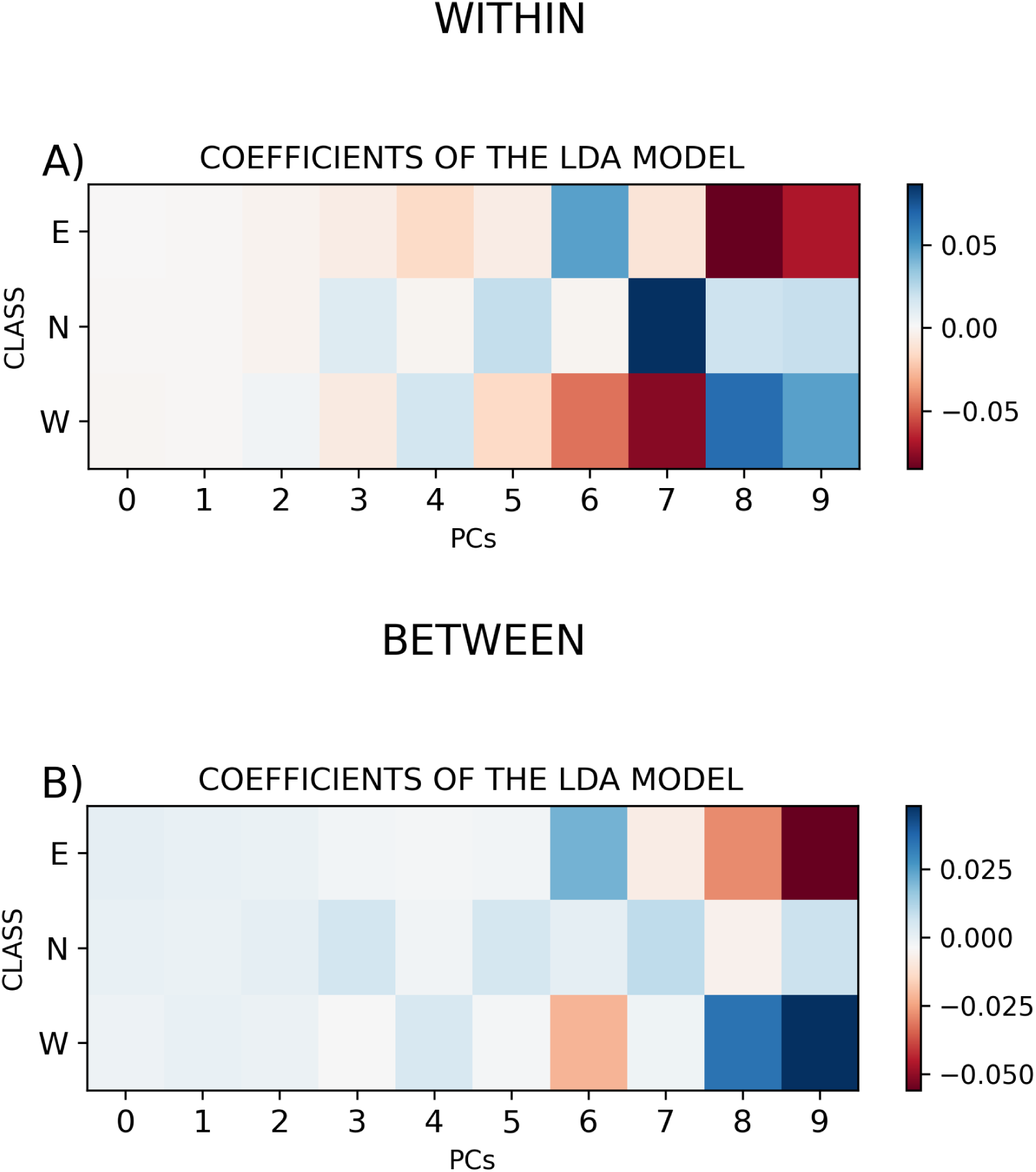
LDA coefficients. The values are the coefficients of the linear combination of the PCs that allows to better discriminate the three classes (E, N, W).

**Figure S11:**
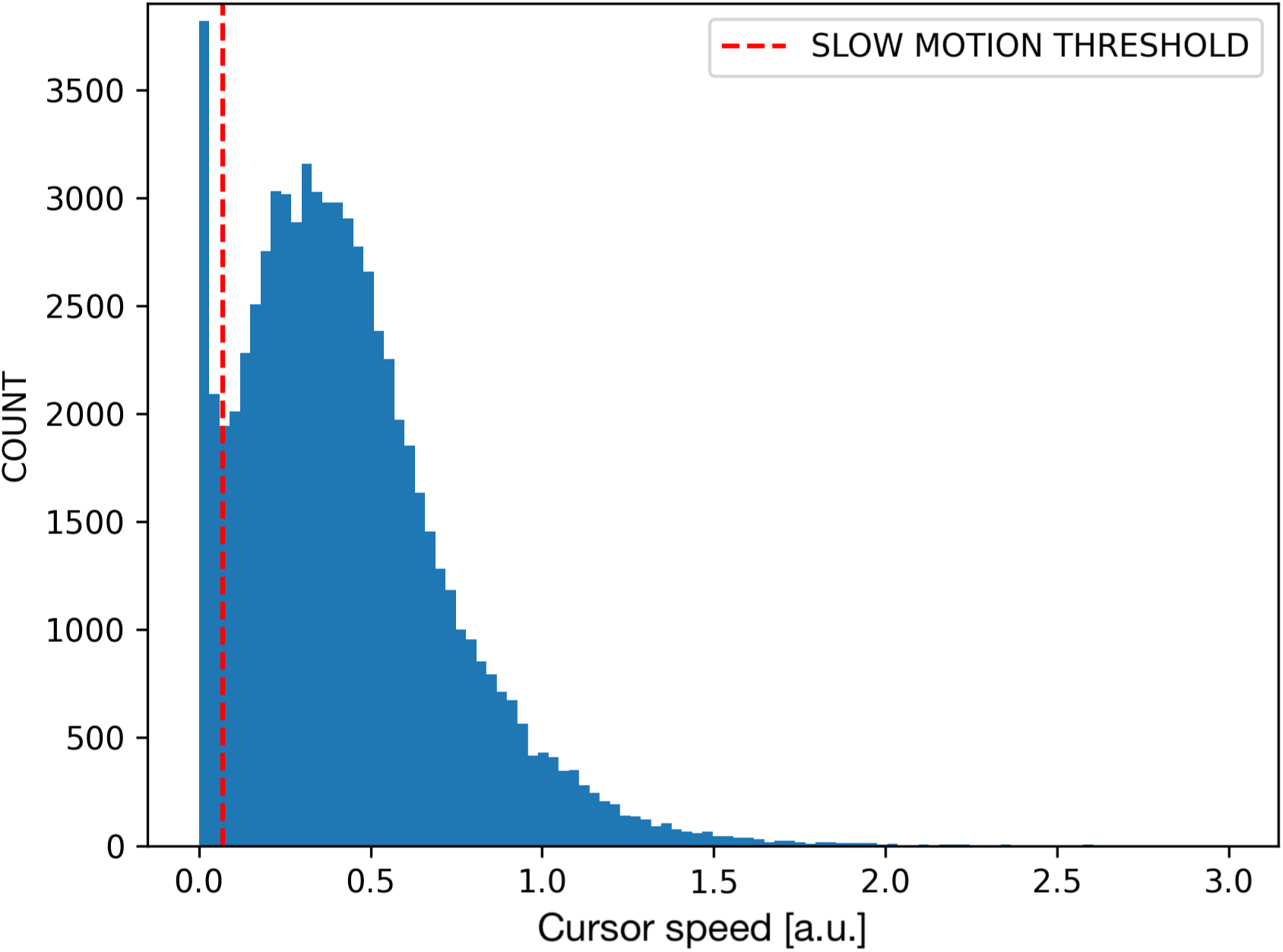
Example of speed threshold for pause detection. The figure shows the speed distribution for cursor movements of a specific subject. The red (dashed) line identifies the first minimum separating the two peaks of the distribution.

## Notes

### Competing Interest Statement

The authors have declared no competing interest.

### Summary of Updates

The paper has been revised to improve methods' clarity.

